# Natural variation in autumn *FLC* levels, rather than epigenetic silencing, aligns vernalization to different climates

**DOI:** 10.1101/2020.04.19.049148

**Authors:** Jo Hepworth, Rea L Antoniou-Kourounioti, Kristina Berggren, Catja Selga, Eleri Tudor, Bryony Yates, Deborah Cox, Barley R Collier Harris, Judith Irwin, Martin Howard, Torbjörn Säll, Svante Holm, Caroline Dean

**Affiliations:** Cell and Developmental Biology, John Innes Centre, Norwich, UK; Computational and Systems Biology, John Innes Centre, Norwich, UK; Department of Natural Sciences, Mid Sweden University, SE-851 70, Sundsvall, Sweden; Department of Biology, Lund University, Lund, SE-223 62, Sweden; Crop Genetics, John Innes Centre, Norwich, UK

## Abstract

Plants monitor temperatures over long timescales to assess seasons and time developmental transitions. In *Arabidopsis thaliana*, winter is registered during vernalization through the temperature-dependent repression and epigenetic silencing of floral repressor *FLOWERING LOCUS C (FLC)*. Natural Arabidopsis accessions show considerable variation in vernalization, however which aspect of the *FLC* repression mechanism is most important for adaptation to different climates is not clear. By analyzing *FLC* silencing in natural variants throughout winter in three field sites, we find that *FLC* starting levels and early phases of silencing are the major variables underlying vernalization response, rather than establishment of epigenetic silencing. This results in an intricate interplay between promotion and delay of flowering to balance survival, and through a post-vernalization effect of *FLC*, reproductive effort via branch production. These data reveal how non-coding *FLC* variation aligns vernalization response to different climatic conditions and year-on-year fluctuations in natural temperature profiles.

**Impact Statement:** Alleles of the major floral repressor vary in their initial expression to underpin the ability of Arabidopsis to survive year-on-year climatic fluctuations.

## Introduction

Developmental transitions in plants are aligned with specific seasons to synchronise with pollinators and optimal climatic conditions (Andrés and Coupland, 2012). A major seasonal cue used to time the transition to flowering is temperature. How temperature affects flowering has been studied genetically in controlled laboratory conditions. However, recent work has shown the importance of analysing this process in natural field conditions (Fig. 1A) (Wilczek et al., 2009; Duncan et al., 2015; Kudoh, 2016; Antoniou-Kourounioti et al., 2018; Hepworth et al., 2018; Rubin et al., 2018; Song et al., 2018; Nagano et al., 2019; Taylor et al., 2019).

**Figure 1.**
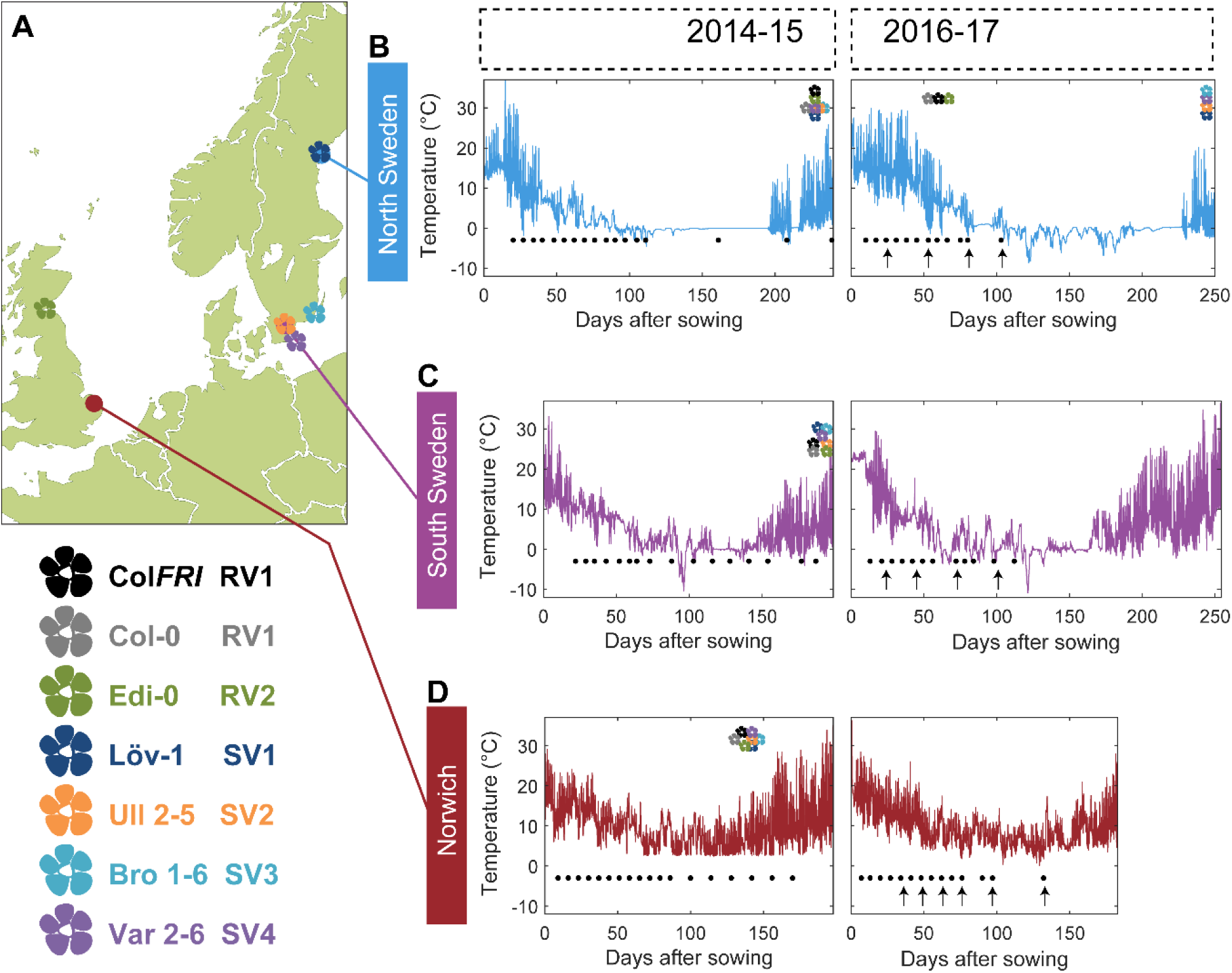
Field experimental setup. **(A)** Map showing locations of field sites (dots) and the origins of five of the accessions (flowers) used in this study. These accessions, with the addition of Col-0, represent the five major and one intermediate (Löv-1) *FLC* haplotypes identified by Li *et al*. (2014). The lab genotype Col *FRI* was also used in this study as a vernalization-requiring reference. **(B-D)** Temperature profiles experienced by plants at the three field sites, North Sweden – Ramsta (B), South Sweden – Ullstorp (C) and Norwich, UK (D) (Source Data 1). Flowers above temperature profile indicate the time of the first bolt of each of the natural accessions and of Col *FRI* (legend at bottom left corner). Black dots below temperature profile indicate the timepoints when plant material was collected for expression analysis. Black arrows below temperature profiles indicate time of transfer to greenhouse with long-day, warm conditions to assess degree of vernalization based on bolting time.

In *Arabidopsis thaliana*, expression of *FLOWERING LOCUS C* (*FLC*), a MADS-box transcription factor that represses flowering, plays a major role in determining the timing of the transition to flowering. *FRIGIDA* (*FRI*) up-regulates *FLC*, establishing an active transcription state at *FLC* chromatin. *FLC* expression is then progressively epigenetically silenced by winter cold, in a process called vernalization (Michaels and Amasino, 1999; Sheldon et al., 1999).

Vernalization represses *FLC* expression through a Polycomb Repressive Complex 2 (PRC2) mechanism that establishes and holds epigenetic memory at *FLC*. This mechanism also involves a temperature-integrating accessory protein, VERNALIZATION INSENSITIVE3 (VIN3) (Sung and Amasino, 2004; De Lucia et al., 2008). Cold-induction of VIN3 leads to formation of a PHD-PRC2 complex, which nucleates Polycomb silencing at an intragenic site within *FLC* (De Lucia et al., 2008). The resulting epigenetic switch and subsequent epigenetic memory is a cis-based mechanism that depends on the local chromatin environment (Angel et al., 2011; Berry et al., 2015), which is influenced by non-coding cis polymorphism at the locus (Lempe et al., 2005; Shindo et al., 2006; Li et al., 2015; Bloomer and Dean, 2017; Sasaki et al., 2018; Qüesta et al., 2020). This non-coding variation defines a small number of major *FLC* haplotypes within the worldwide *Arabidopsis thaliana* population, which confer different vernalization responses and appear to have been maintained in the *A. thaliana* population due to their contributions to life history diversity (Shindo et al., 2006; Li et al., 2014).

Previous field work in Sweden and Norwich, UK, with the vernalization reference genotype Col *FRI*^SF2^, and mutants in *VIN3* (*vin3-4*) had shown that initial *FLC* down-regulation requires cold temperature, whereas the epigenetic silencing requires the absence of daily temperatures above 15 °C (Hepworth et al., 2018). In different climates the autumn conditions greatly influenced when epigenetic silencing initiated (Hepworth et al., 2018). These data were used to develop a mathematical model of vernalization that can predict *FLC* silencing in natural field conditions (Antoniou-Kourounioti et al., 2018; Hepworth et al., 2018). However, the importance of cis *FLC* polymorphism in delivering the different phases of *FLC* silencing in different climates was unknown. Here, we exploit our field studies in three climatically very distinct locations, over multiple years with a high degree of climate variation, to investigate genotype versus environment interaction of *FLC* cis polymorphism in vernalization response (Fig. 1).

Our results demonstrate how the major *FLC* haplotypes, differing only through non-coding variation, have different starting *FLC* levels and rates of response to autumn cold, but show remarkably similar epigenetic silencing rates in winter. This generates a uniform vernalisation response in most years across haplotypes and climates in the field. However, in capturing an unusual year, our experiments also reveal effects of the haplotypes on reproductive success through branching and higher silique number. By studying gene expression across years and climates, we have been able to dissect how non-coding cis variation effects are important in the adaptation of vernalization response to natural fluctuating environments.

## Results

### Field experiments

Li *et al*. (2014) identified 20 haplotypes representing the major allelic variation at the *FLC* locus across a worldwide panel of more than a thousand Arabidopsis accessions. Although the variation characterising these haplotypes is entirely due to non-coding or synonymous single-nucleotide polymorphism, these haplotypes conferred different responses to vernalization in laboratory conditions (Li et al., 2014). To investigate their function in field conditions, we selected accessions to represent each of the five most populous haplotypes, in total representing more than 60% of tested accessions, as well as a further accession, Löv-1, for which there is evidence of local adaptation to the climate in the region of our North Sweden field site (Duncan et al., 2015; Qüesta et al., 2020). To compare these alleles in a common genetic background, we exploited extant and developed new Near Isogenic Lines (NILs) in which the *FLC* allele from each accession had been repeatedly backcrossed to Col *FRI*^SF2^ (“Col FRI”; Duncan et al., 2015; Li et al., 2015). These genotypes were tested across two years and three field sites, with the exception of three NIL lines which were synthesised during the experiments (Fig. 1A, 2B-J). The three field sites in Norwich, UK, in Ullstorp, Sweden (“South Sweden”) and in Ramsta, Sweden (“North Sweden”) were chosen to represent different climates, and are close to the source sites of several of the tested accessions; Vår2-6 and Ull2-5 in Skåne, near or at Ullstorp; Löv-1 near Ramsta (Fig. 1A). In the second year of experimentation, we also included the *vin3-1* mutant in the Col *FRI* background. The experiments ran from August 2014 until spring 2015 and again from August 2016 to the spring of 2017. In the first year, two plantings were performed in North Sweden, two weeks apart. The temperatures that the plants experienced are shown in Fig. 1A. We measured the levels of spliced and unspliced *FLC*, and mRNA levels of the key cold-responsive input to *FLC, VIN3*, to follow the progress of vernalization in the field. The transition to flowering (bolting) was assessed both in the field and by transfers to warm inductive conditions.

### Natural variation in different phases of *FLC* silencing in the field

Across all the genotypes we tested and all seven field experiments, as expected, *FLC* expression reduced over weeks in response to autumn and winter temperatures, whereas *VIN3* was upregulated. Previously, we had noted that in 2014-5 in Norwich, substantial *VIN3* upregulation did not occur until ∼65 days after sowing, although temperature conditions were suitable for VIN3-independent *FLC* downregulation for most of this time (Hepworth et al., 2018). In the following field season, this pattern occurred again, with *VIN3* upregulation delayed until 48 days after sowing (Antoniou-Kourounioti et al., 2018). For the Col *FRI* reference, we had found that we could fit two separate exponential decay curves to *FLC*; the first for the initial, slow, VIN3-independent phase and the second for the faster, VIN3-dependent phase of the downregulation (Fig. 2A; Hepworth et al., 2018). Thus three features of the *FLC* profile contribute to the level of *FLC* at any time: firstly the ‘starting level’ of *FLC* before vernalization, secondly the rate of downregulation in the initial VIN3-independent phase, and thirdly the rate of VIN3-dependent downregulation (Fig. 2A). This pattern was consistent across the accessions and NILs, allowing us to investigate the effect of natural variation on different aspects of *FLC* regulation. The time of upregulation of *VIN3*, and thus the time of switching from the VIN3-independent to the VIN3-dependent shutdown, also affects *FLC* levels, but this was very similar between genotypes at the same site.

**Figure 2.**
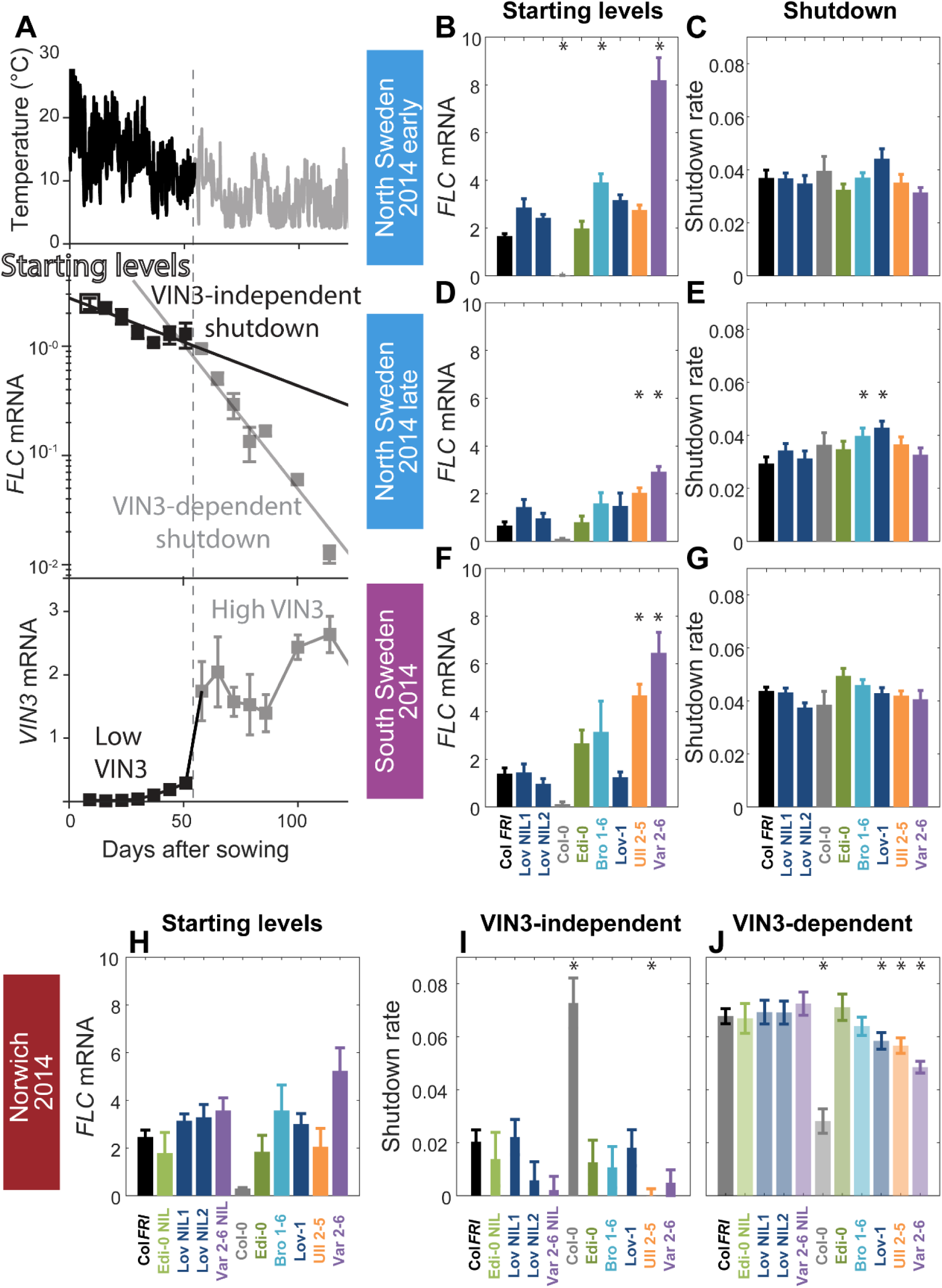
Downregulation in 2014/15 in Norwich, North Sweden (2 plantings) and South Sweden for all NILS and accessions. **(A)** Experimental data for Col *FRI* in Norwich 2014-5, showing the temperature profile (top), *FLC* (middle) and *VIN3* (bottom) expression. Different shades indicate the separation of the VIN3-dependent (grey) and -independent (black) phases of *FLC* silencing (Hepworth et al., 2018) and equivalent times in *VIN3* and temperature profiles. Expression data was normalised to the control sample for 2014-5 (see Methods). N=6 except where samples were lost to death or degradation (see Methods and Source Data 2). The initial measurement in the field (Starting Levels), the rate of downregulation before induction of *VIN3* expression (VIN3-independent, estimated from the slope of the fitted line) and the rate of downregulation after *VIN3* induction (VIN3-dependent) are the three features that were analysed and compared for each genotype and treatment. Error bars show standard error of the mean (s.e.m). **(B-J)** *FLC* downregulation analysed as level at first time point (Starting levels), and rate of downregulation (Slope) for North (B-E) and South Sweden (F-G), or rate of downregulation before (Slope, dark bars) and after (Slope, translucent bars) *VIN3* induction for Norwich (H-J). Features of genotypes that are significantly different to the reference line Col *FRI* are indicated by *. p-values for all comparisons are given in Supplementary file 1. Rates of downregulation are given in units of “a.u. per day”, where the arbitrary units (a.u.) correspond to the normalised concentration of *FLC* mRNA. *VIN3* induction started at ∼58 days in Norwich (Fig. S2-2, S2-3). Expression data was normalised to the control sample for 2014-5 (see Methods). Error bars show s.e.

In Norwich 2014-15, the ‘Rapid Vernalizing’ (RV) *FLC* accession Edi-0 (haplotype group RV1) behaved very similarly to Col *FRI* (RV2). In comparison, the ‘Slow Vernalizing’ (SV) haplotypes (Löv-1, Ull2-5, Bro1-6 and Var2-6) show higher levels of *FLC* throughout the winter (Fig. 2, S2-1). Ull2-5 started with similar levels of *FLC* as Col *FRI*, but slower rates of downregulation in both the VIN3-independent and VIN3-dependent phases generated higher levels of *FLC*. For both Var2-6 and its NIL, an apparently slower VIN3-independent rate of downregulation contributed to its raised *FLC*, indicating that the cis variation at *FLC* was responsible for this difference. The VIN3-dependent phase was also slower in Var2-6 and Lov-1 (Fig. 2), but not in their NILs, suggesting that this effect is not governed by the *FLC* locus.

In Sweden 2014-5, at both sites the VIN3-independent and -dependent phases occurred concurrently (Fig. 2, S2-1; Antoniou-Kourounioti et al., 2018). There was little variation observed in the overall rate of *FLC* downregulation between genotypes, though *VIN3* induction was more variable (Fig. S2-2). Instead, most of the natural variation in *FLC* levels throughout winter in Sweden was generated by differences in the early expression level. In North Sweden again both RV accessions had similar starting levels whereas the SV Swedish accessions were higher to different degrees. However, in South Sweden Löv-1 starts similarly to Col *FRI*.

For the 2016-7 experiment, we sowed plants in North Sweden and Norwich two weeks earlier and South Sweden three weeks earlier than for the 2014-5 season (Fig. 1A, 3, S3-1, S3-2). In North Sweden this was followed by a warm autumn and produced a delay in *VIN3* induction similar to that seen in both years in Norwich. However, the first stage of shutdown was still more rapid in North Sweden than Norwich, despite higher average temperatures in North Sweden (Fig. 1A), so that both VIN3-independent and VIN3-dependent phases had similar rates, resulting in the appearance of a single decline. As in the previous experiment in Sweden, these rates of downregulation were generally similar between different genotypes, with higher *FLC* levels in SV accessions and the Var NIL due to higher starting levels (Fig. 3, S3-1).

**Figure 3.**
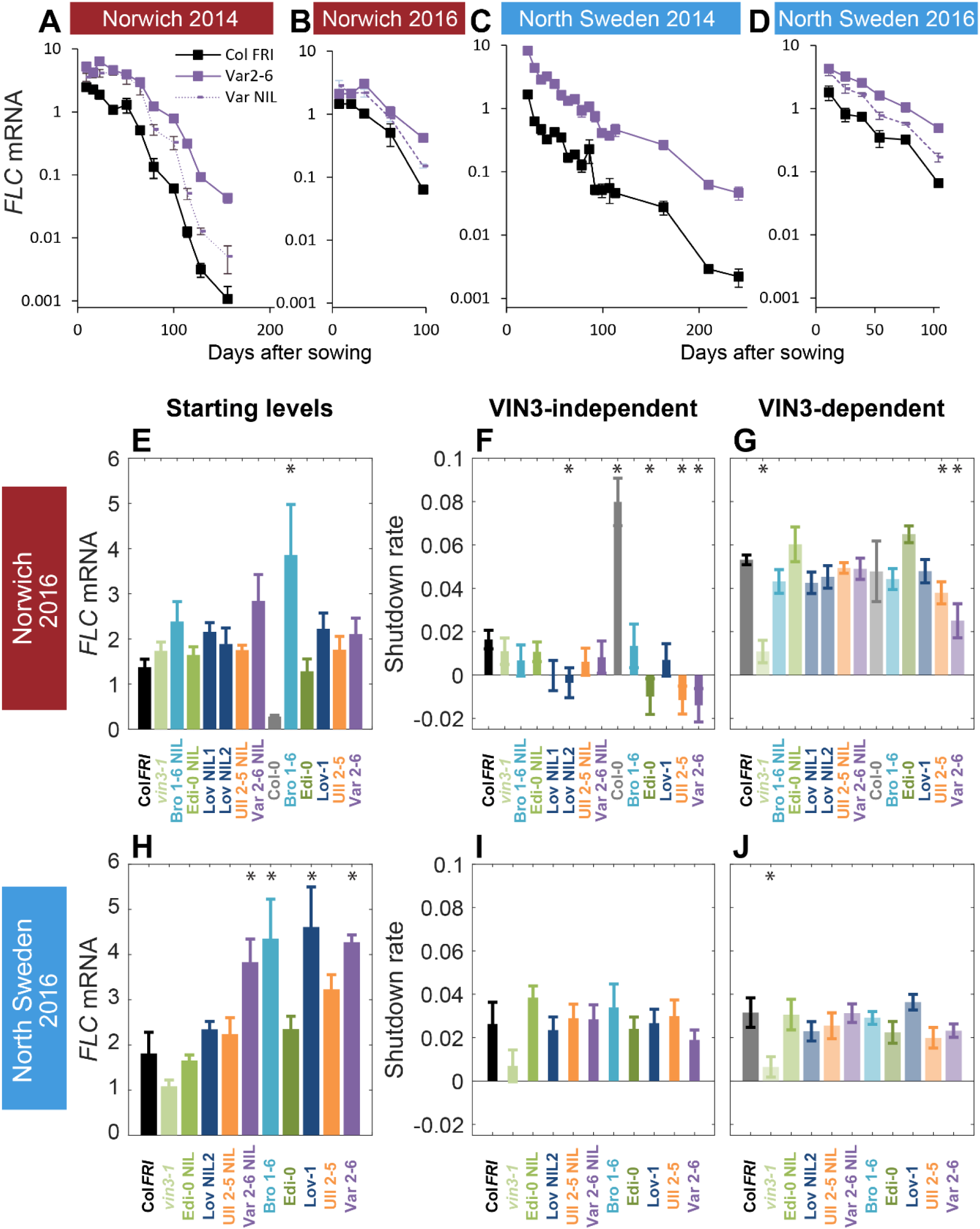
Downregulation in 2016 in Norwich and North Sweden for NILS and accessions show similar patterns of response to the first year. (**A-D**), *FLC* downregulation in Col *FRI*, Var2-6 and the Var NIL, as measured for Norwich and North Sweden in the winters of 2014-5 and 2016-7. (**E-J**), *FLC* downregulation as starting level and VIN3-independent and dependent rates. Features of genotypes that are significantly different to the reference line Col *FRI* are indicated by *. p-values for all comparisons are given in Supplementary file 1. Expression data was normalised to the corresponding control sample (2016-7, see Methods). N=6 except where samples were lost to death or degradation (see Methods and Source Data 3). Rates of downregulation are given in units of “a.u. per day”, where the arbitrary units (a.u.) correspond to the normalised concentration of *FLC* mRNA. Error bars of bar plots show s.e., of line graphs show s.e.m.

In Norwich the patterns seemed to be largely repeated, though our statistical power was lower in the 2016-7 experiment due to fewer timepoints. Bro1-6, Lov-1, Var 2-6 and their NILs appeared to have higher starting *FLC* levels, though of these only Bro1-6 was significant in our analysis. Ull2-5, Var2-6, Edi-0 and Löv NIL2 showed slower VIN3-independent downregulation and Ull2-5 and Var2-6 also again showed slower VIN3-dependent downregulation (Fig. 3). Other than this slower VIN3-independent rate in Norwich in Edi-0, the RV alleles behaved similarly to each other again in 2016-7.

The slower rate of the later phase in Var2-6 was consistent across years in Norwich (Fig. 2, 3). Unlike the other accessions, this change in the VIN3-dependent phase in Var2-6 was consistently mirrored in lower levels of *VIN3* (Fig. S2-2, S3-1). The circadian clock is an important regulator of *VIN3* (Antoniou-Kourounioti et al., 2018; Hepworth et al., 2018). When sampled over 48 hours in the Norwich field experiment (Fig. S3-3), *VIN3* expression in Var2-6 is much lower compared to our previous results from Col *FRI* (Fig. S3-3B, D; Antoniou-Kourounioti et al., 2018), as is the expression of circadian clock component *CIRCADIAN CLOCK ASSOCIATED1*, the protein of which binds to the *VIN3* promoter (*CCA1*; Fig. S3-3; Nagel et al., 2015). Therefore, variation in circadian regulation may underlie some of the difference in *FLC* regulation in Var2-6.

### Chromatin modifiers control *FLC* regulation in the field

In Norwich, our contained site allowed us to investigate mutants in genes known to affect *FLC* levels before cold or in response to cold to look at what trans factors may provide temperature information to the different phases of shutdown (Fig. 4).

**Figure 4.**
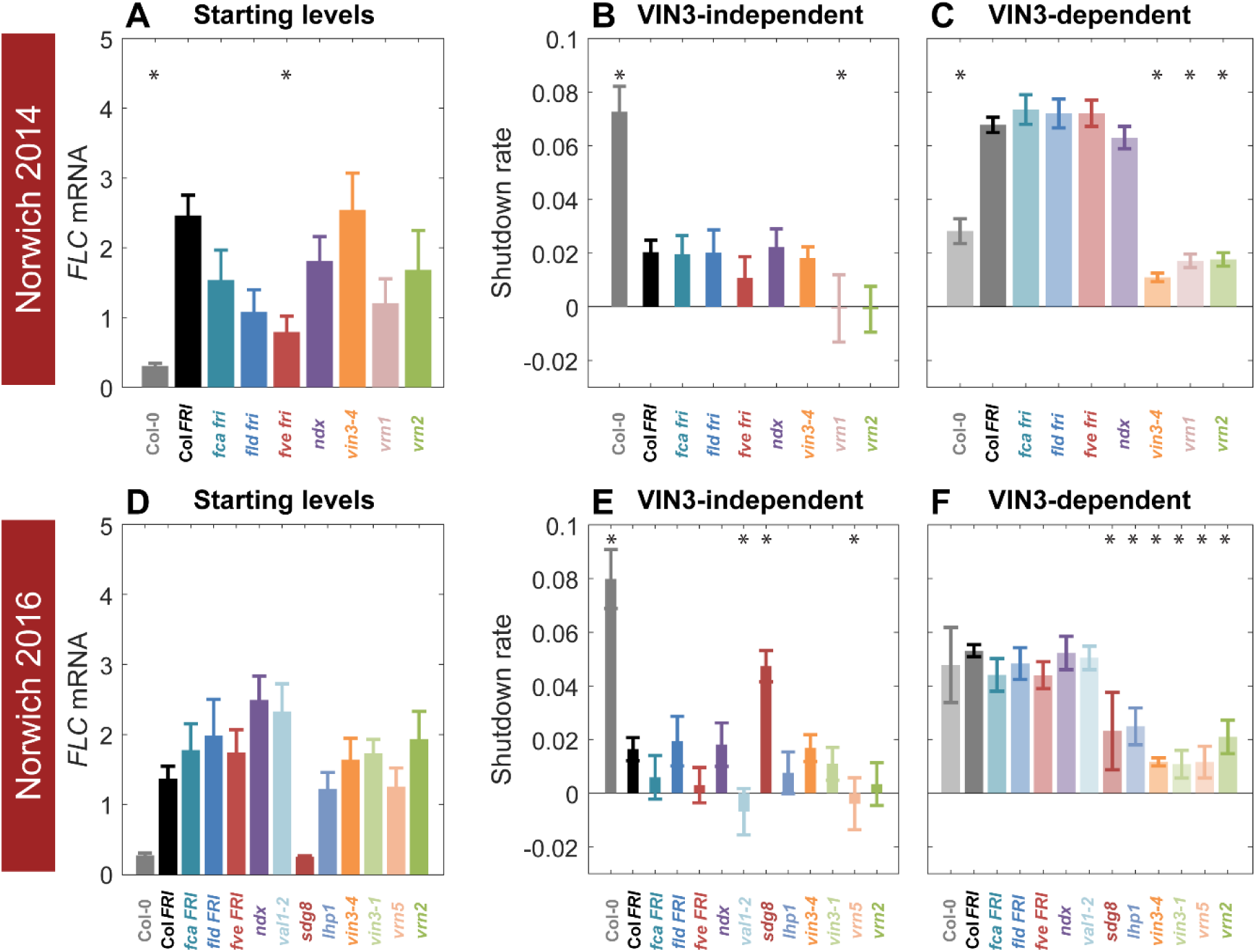
Starting levels and rates of downregulation of *FLC* in mutants and transgenics in field conditions in Norwich, UK. (**A-F**), *FLC* downregulation analysed as level at first time point (Starting levels, A, D), rate of downregulation before induction of *VIN3* expression (Slope, dark bars, B, E) and rate of downregulation after *VIN3* induction (Slope, translucent bars, C, F). Features of genotypes that are significantly different to the reference line Col *FRI* are indicated by *. p-values for all comparisons are given in Supplementary file 1. Rates of downregulation are given in units of “a.u. per day”, where the arbitrary units (a.u.) correspond to the normalised concentration of *FLC* mRNA. *VIN3* induction started at: Norwich 2014, ∼58 days, see Fig. S2-2; Norwich 2016, ∼48 days, South Sweden 2016, ∼35 days, North Sweden, ∼46 days, see Fig. S3-1). All mutants are in the Col *FRI* background unless otherwise stated. Expression data was normalised to the corresponding control sample (for 2014-5 or 2016-7, see Methods). N=6 except where samples were lost to death or degradation (see Methods and Source Data 2 and 3). Error bars show s.e.

As previously reported (Hepworth et al., 2018), mutants in VIN3 (*vin3-4, vin3-1*) did not show the increase in downregulation rate that marks the later epigenetic phase of silencing (Fig. 4C, F). This effect was also seen in other epigenetic memory mutants; VERNALIZATION1 (*vrn1-4*); the PRC2 component VRN2 (*vrn2-1*); the VIN3-related protein VRN5 (*vrn5-8*); and LIKE HETEROCHROMATIN1 (*lhp1-3*), in accordance with this phase representing epigenetic silencing. Loss of the H3K36 methyltransferase SET DOMAIN GROUP8 (*sdg8*) generated the same effect, correlated with its delayed upregulation of VIN3 (Kim et al., 2010; Finnegan et al., 2011).

In the initial, VIN3-independent silencing phase, both Col *fri* and *sdg8 FRI* were hyperresponsive, consistent with SDG8’s role with FRI in the establishment of high *FLC* transcription (Hyun et al., 2017). Mutants in the ‘autonomous flowering’ pathway, which upregulate *FLC* expression in the absence of *FRI*, had no significant effect on VIN3-independent silencing, but behaved mostly like Col *FRI*, although *fve-3* reduced the rate of downregulation non-significantly in both years. The mutant with the most dramatically reduced VIN3-independent response was the B3-binding transcription factor VAL1 (VP1/ABI3-LIKE 1), required for PRC2 action at *FLC* (Fig. 4E; Qüesta et al., 2016; Yuan et al., 2016), followed closely by the *vrn* mutants, *vrn1* and *vrn5*. Conversely, the *lhp1-3* and *vin3* mutants clearly show no impairment in the early phase of downregulation. Overall, it seems that epigenetic silencing components are required for both phases of downregulation in the field.

### Initial levels of *FLC* are the major variables in vernalization response

To identify the major variable aligning vernalization response to different climates, we estimated the coefficient of variation for the rates of shutdown and starting levels for all the natural accessions, NILs, and *vin3-1* where available (from Fig. 2, 3, S2-1, S3-1, Supplementary Table 3). In Sweden the starting levels are significantly more variable than the slopes (p-value=2.8 · 10^−8^ for the first year, Fig. 5A) but in Norwich the VIN3-independent shutdown rate is more variable than the starting levels and VIN3-dependent rate (p-value=2.1 · 10^−5^ for 2014, p-value=5.0 · 10^−4^ for 2016, Fig. 5B). Combining Norwich and Sweden data, the early shutdown rate is again most variable (p-value=1.9 · 10^−10^ for 2014, p-value =2.0 · 10^−5^ for 2016, Fig. 5C). On the other hand, there was no significant difference in the variability of the starting levels (Fig. 5D) between the different field sites and years, and similarly for the shutdown rates (Fig. 5E-G). What we describe as the starting level was measured after some days in the field, so it is not equivalent to a non-vernalized control. Some *FLC* shutdown, most likely VIN3-independent, will have occurred at that time. Therefore, it is likely that the combination of these two determinants of *FLC* levels early in the field (starting levels, VIN3-independent shutdown) provides most of the potentially adaptive variation.

**Figure 5.**
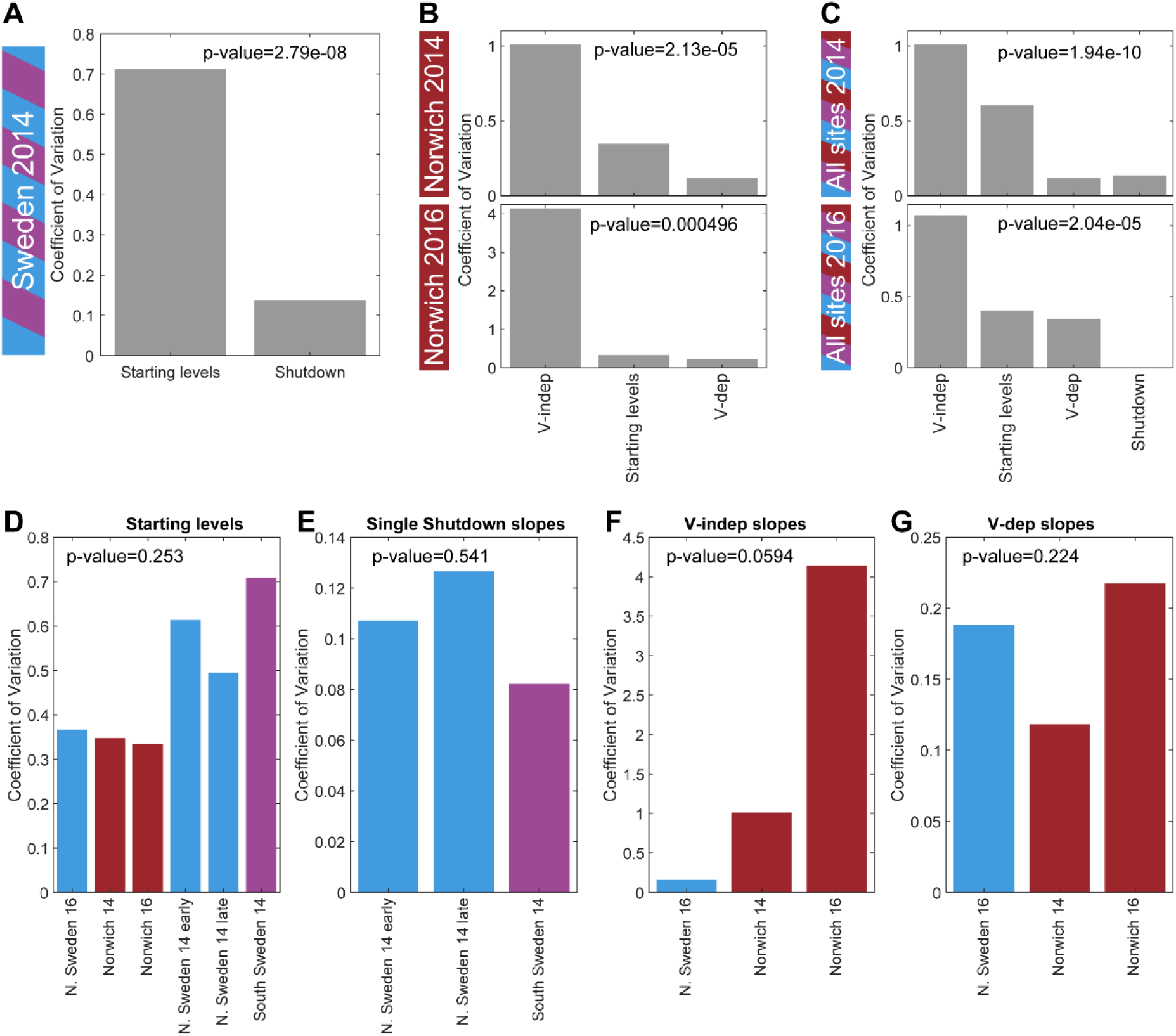
Causes of natural variation in *FLC* levels across sites and years. **(A)** The coefficient of variation for the rates of shutdown and for the starting levels in all Sweden experiments in the first year. **(B)** Similarly in Norwich 2014 (top) and 2016 (bottom) but separately for the VIN3-independent (V-indep) and VIN3-dependent (V-dep) shutdown rates. **(C)** Comparison of the variability of the starting levels and the shutdown rates, separating V-dep and V-indep where appropriate, combining data from all sites in 2014 (top) and 2016 (bottom). “Shutdown” refers to the combined V-dep/V-indep shutdown rate that was fitted in Sweden 2014, and so is not present in the 2016 results. **(D)** The coefficients of variation of the starting levels for each site/year. **(E)** The coefficients of variation of the single shutdown rates for the different plantings and sites in Sweden in 2014. **(F-G)** Similarly, for Sweden 2016 and Norwich in both years, separating the V-indep rates (F) and V-dep (G). Data from Source Data 2 and 3.

### Vernalization is saturated before midwinter in the field

Having characterised the variation in *FLC* response to field conditions, we investigated the effect of this variation on the floral transition in the field.

In the 2014-5 season, accessions and NILs bolted quite synchronously in Norwich and Sweden, with only the Bro1-6 accession in Norwich showing much relative delay in the floral transition (Figure S6-1 and S6-2). These results suggested that the vernalization effect on *FLC* is normally saturated across accessions and climates in the field. To test this hypothesis, in 2016-7 we removed plants from field conditions to heated, long-day conditions to induce flowering and scored the length of time until floral buds were visible at the shoot apex (bolting). There was wide variation in the delay to bolting between and within the genotypes for plants transferred to the warm greenhouses early in autumn, at all sites, indicating that vernalization requirement had not been saturated at this point (Fig. 6A, C, E). However, as winter progressed, time to bolt reduced and became more uniform for plants within and between each genotype (Fig. 6B, D, F), although vernalization saturated at different rates in different genotypes (Fig. S6-3, S6-4). All accessions and NILs bolted broadly synchronously after removal on 21^st^ December in Norwich, in South Sweden by 17^th^ December, and in North Sweden accessions were almost synchronous by the date of the final transfer, 24^th^ November (Fig. 6, S6-3, S6-4), in line with previous findings for North Sweden (Duncan et al., 2015). Therefore, in current climates almost all Arabidopsis plants have probably saturated their requirement for vernalization well before midwinter.

**Figure 6:**
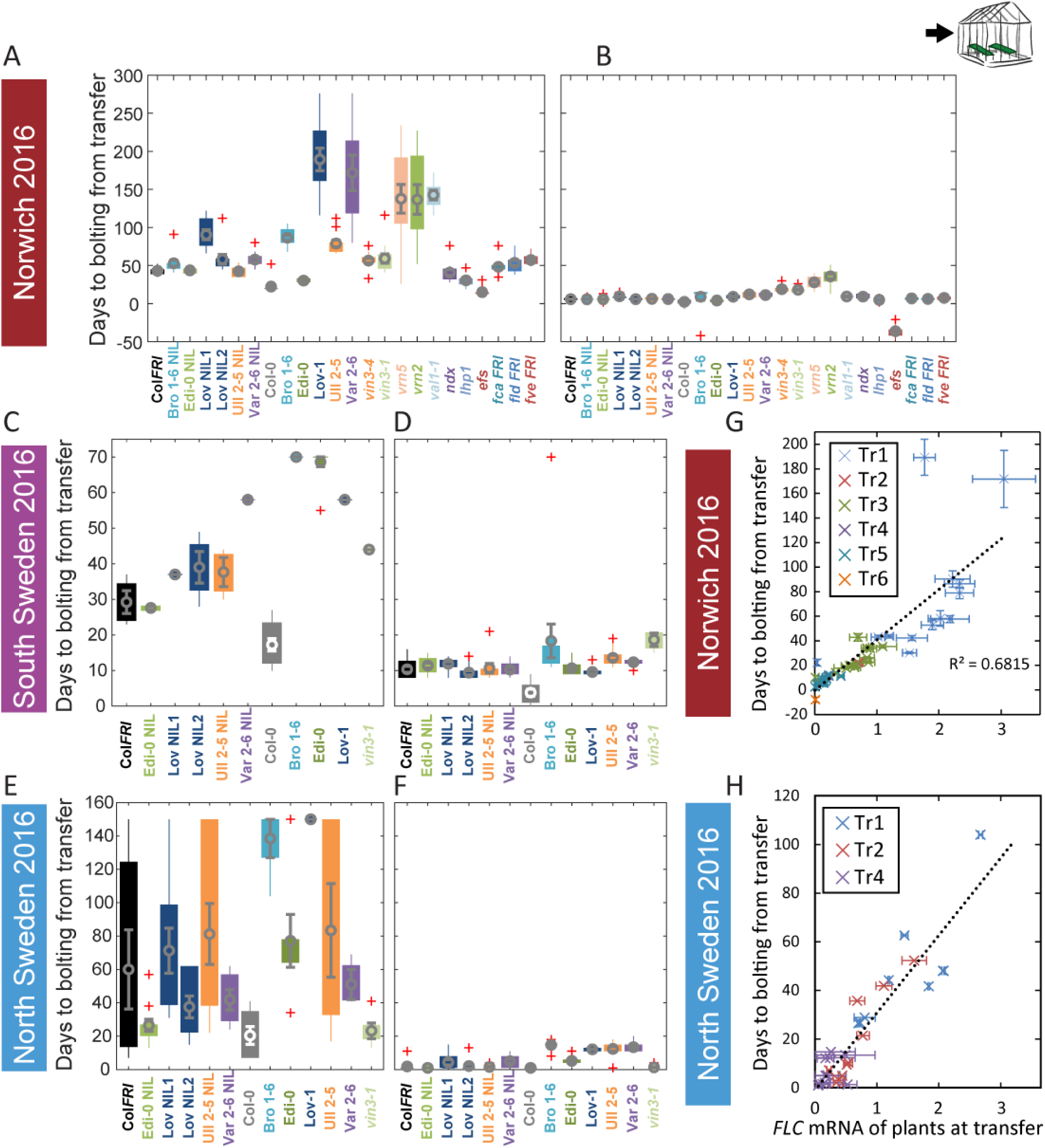
Vernalization requirement for *FLC* downregulation is saturated in natural winters. **(A-F)** Bolting time for accessions and NILs after transfer to floral-induction conditions from ‘natural’ winter 2016-7, in **(A)** Norwich 21/10/16 **(B)** Norwich 21/12/16 **(C)** South Sweden 01/10/2016 **(D)** South Sweden 17/12/16 **(E)** North Sweden 06/09/2016 **(F)** North Sweden 24/11/2016. Plants that did not flower within 70 days (C, D) or 205 days (E, F) not shown. **(G-H)** North Sweden 2016 transfers for accessions and NILs, **(G)** mean time to bolting after transfer to floral-inductive conditions plotted against mean *FLC* expression per genotype at transfer, Norwich 2016-17, R^2^= 0.68, p<0.001. **(H)** mean time to bolting after transfer to floral-inductive conditions plotted against mean *FLC* expression per genotype at transfer, North Sweden 2016-17, genotypes that did not bolt within 205 days not shown, R^2^= 0.85, p<0.001. N=12 plants except where plants died or (E, H) did not bolt within 205 days (Source Data 5). Error bars show s.e.m.

### The same expression levels from different *FLC* alleles predict different flowering transition responses in transfer experiments

Across genotypes, in Norwich and North Sweden 2016-7, the time to the floral transition correlated closely with the *FLC* expression at the time of transfer to floral inductive conditions, as expected (North Sweden, R^2^ = 0.85, p<0.001, Norwich, R^2^= 0.68, p<0.001, linear regression Fig. 6G, H). Accessions and NILs with high starting levels and slower downregulation rates generally bolted later and took longer to saturate their vernalization requirement.

However, each accession differed in the relationship between *FLC* levels at transfer and subsequent bolting time (Fig. S6-5, S6-6). This is partly due to variation at the many other genes that regulate the floral transition, but it was also observed among the NILs, suggesting that this relationship is also controlled by *cis* variation at *FLC*. For example, for Löv-1 and the Löv NIL1, the time to bolt for a given level of *FLC* at transfer is longer than for Col *FRI*, but for Edi-0, it is shorter (Fig. S6-5, S6-6).

This analysis allowed us to extract a further feature of *FLC* regulation, the relationship of *FLC* levels at the time of transfer to the floral transition time. We named this feature the *FLC*-post-vern value (*m* in Table 1). An allele with a higher *m* value suggests that at a given level of *FLC*, this allele results in later flowering than an allele with a lower *m* value would at the same expression value of *FLC* at the time of transfer. We calculated this value from both the Norwich and the North Sweden transfers. There was substantial variation between the different glasshouses in the estimates for the accessions (Table 1). However, for the lines in the common Col *FRI* background the estimates generally correlated well, with the exception of the Ull NIL. Therefore, within the Col *FRI* background, we can quantify our previous observations and use the *FLC* levels to roughly predict the expected bolting time.

**Table 1.** Linear regression relationship between bolting time and *FLC* mRNA expression, as shown in Fig. S6-5 and S6-6, where *days to bolting = m[FLC mRNA] + c*, and *m* is the ‘*FLC-* post-vern’ value and *c* is a fitted constant relating to non-FLC-mediated bolting delay. NA – estimate only based on two data points, so no standard error is calculable. n.d – no data.

| Genotype | Norwich<br>*m* | Norwich<br>Std. error | Norwich<br>p value | North<br>Sweden<br>*m* | North<br>Sweden<br>Std.<br>error | North<br>Sweden<br>p value | Average<br>*m* |
| --- | --- | --- | --- | --- | --- | --- | --- |
| Col *FRI* | 36.5 | 5.5 | 0.00704 | 39.0 | 17.6 | 0.269 | 37.7765 |
| *vin3-1 FRI* | 51.0 | 3.9 | 0.0486 | 67.3 | 27.6 | 0.247 | 59.14 |
| Bro NIL | 36.1 | 0.33 | 0.00583 | n.d. | n.d. | n.d. | 36.1209 |
| Edi NIL | 33.5 | 2.7 | 0.0507 | 39.8 | 1.08 | 0.0173 | 36.68155 |
| Löv NIL1 | 40.7 | 5.08 | 0.0791 | n.d. | n.d. | n.d. | 40.715 |
| Löv NIL2 | 27.4 | 0.80 | 0.0186 | 23.8 | 0.507 | 0.0136 | 25.5717 |
| Ull NIL | 24.8 | 0.98 | 0.0252 | 42.1 | 9.27 | 0.138 | 33.43305 |
| Var NIL | 25.7 | 0.89 | 0.0221 | 22.3 | 1.60 | 0.0456 | 23.9947 |
| Col-0 | 486 | 155 | 0.197 | n.d. | n.d. | n.d. | 486.436 |
| Bro1-6 | 27.0 | 0.31 | 0.00736 | 37.3 | 2.08 | 0.0355 | 32.1373 |
| Edi-0 | 16.4 | 3.2 | 0.124 | 48.3 | 9.15 | 0.119 | 32.395 |
| Löv-1 | 113 | 19 | 0.11 | n.d. | n.d. | n.d. | 113.08 |
| Ull2-5 | 31.4 | 1.2 | 0.0238 | 16.9 | 8.54 | 0.297 | 24.1725 |
| Var2-6 | 63.1 | 6.9 | 0.0698 | 35.0 | NA | NA | 49.0475 |

### High *FLC* reduces precocious flowering in a warm autumn

At all sites in the 2014-15 season, and in Norwich in 2016-17, none of the natural accessions or NILs flowered in the field until after mid-winter (Fig. S6-1, S6-2). Therefore, in at most sites in most years the variation in *FLC* levels in autumn that we had observed had no phenotypic consequence. However in North Sweden winter 2016-17 many of the plants with the Col background (NILs, Col *FRI*, and *vin3-1 FRI*) transitioned to flowering early, by the 18^th^ November (65 days after sowing), before winter and snowfall (Fig. 7A). Precocious bolting was rare in the SV accessions and was much reduced even in the RV accession Edi-0. Over all the genotypes, the percentage of plants transitioning to flowering before winter negatively correlated with genotype *FLC* expression on 5^th^ October, one day after the first recorded bolting for plants in the field (p<0.001, GLM for binomial data, Fig. 7B). Within the Col *FRI*^SF2^ background, the SV Var and Löv *FLC* alleles, the NILs with the highest *FLC* levels during autumn (Fig. S3-1), substantially reduced precocious bolting (p=0.005, binomial proportions test, Fig. 7A).

**Figure 7:**
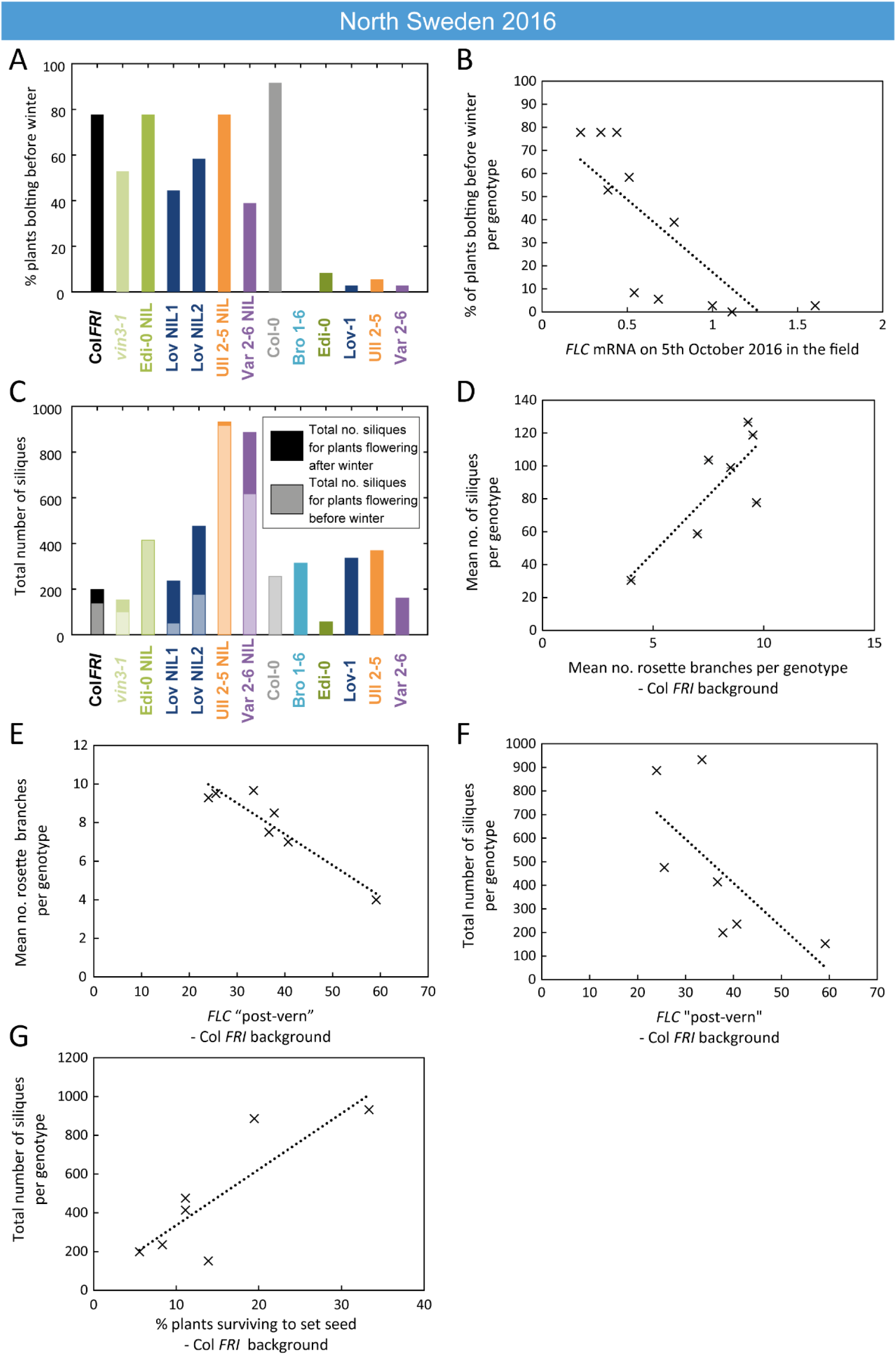
High *FLC* genotypes reduce precocious bolting in North Sweden in warm years. **(A)** Percentage of plants bolting before winter in the North Sweden 2016 experiment by genotype. Plants in the field are less likely to flower precociously before winter (18^th^ November 2016) if they are accessions from more northerly latitudes or, to a lesser degree, if they are *FLC* introgression lines from these SV haplotypes. **(B)** The percentage of plants transitioning to flowering before winter per genotype negatively correlated with *FLC* expression (normalised to control) on 5^th^ October (R^2^=0.59, p = 0.0058). **(C)** Total number of siliques produced per genotype, showing contribution from plants that bolted before winter and plants that bolted after. Within the Col *FRI* genetic background. no overall penalty in average silique number for surviving plants bolting before versus after winter (92 and 77 per plant respectively, not significant in Mann-Whitney U test). **(D)** Mean silique production in plants surviving to set seed positively correlated to their mean rosette branch production for Col *FRI* genetic background genotypes (NILs and *vin3-4*) (R^2^ = 0.56, p-value = 0.002). **(E)** Rosette branch production of Col *FRI* genotypes surviving to set seed is strongly negatively correlated with the *FLC* post-vern value for that genotype as from Table 1 (R^2^ = 0.86, p-value < 0.002). **(F)** Total number of siliques produced by Col *FRI* background genotypes plotted against *FLC* post-vern, linear regression for post-vern effect alone, R^2^ = 0.35, p-value = 0.1. **(G)** Total number of siliques produced by Col *FRI* background genotypes plotted against percentage survival of that genotype to point of seed set, linear regression for survival effect alone, R^2^ = 0.64, p-value = 0.019. N=36 plants sown (A-C), n for surviving plants (D-G) varies per genotype, see Source Data 6.

Across all genotypes, plants that bolted precociously were not observed to set seed before winter and were less likely to survive the winter (42% of bolting plants survived, whereas 67% of non-bolting plants survived to spring, p<0.001, binomial proportions test, Fig. S7-1A). Therefore, high *FLC* levels in October correlated with higher survival to seed set in the field (p<0.003, GLM for binomial data, Fig. S7-1B).

### Variation at *FLC* affects fitness in the field through branching and silique number

Although survival is critical for fitness, subsequently the number and viability of the seeds set by the plant determines reproductive success. We confirmed the promotive effect of vernalization on branching in *A. thaliana* in our core accessions and NILs in controlled conditions (Fig. S7-2, S7-3) and noted that NILs bearing different *FLC* alleles showed different rates of branch production in response to differing lengths of vernalization (Fig. S7-3C). Moreover, in the 2016 Norwich transfer experiments, plants that were transferred before the vernalization requirement was saturated produced lower and more variable amounts of seed (Fig. S7-4).

To investigate the role of *FLC* specifically on silique numbers, we looked at the behaviour of the genotypes within the Col *FRI* genetic background. In these plants, there was no overall penalty in average silique number for plants that bolted before winter compared to those that transitioned afterwards, provided that they survived (Fig. 7C, S7-1D).

Silique production occurred only after winter, after saturation of vernalisation requirement (Fig. 6F) and yet there was large variation in branching and silique production within the Col *FRI* background plants. We asked whether the likelihood of *FLC* reactivation after winter might relate to fitness. To estimate this, we tested the relationship between *FLC* and bolting time in the warm, the ‘*FLC-*post-vern value’ as derived from Fig. S6-5 and Fig S6-6 (Table 1), however, this did not seem to affect survival to seed set or date of bolting of the survivors (Fig. S7-1E, F). Instead, within the Col *FRI* background, the average silique set amongst survivors correlated with rosette branching (Fig. 7D, Fig. S7-1G) and branch number was strongly negatively correlated with *FLC*-post-vern value (Fig. 7E). Between them, survival to seed set and post-vern value explained a large part of the variation in total silique set per Col *FRI* genotype (linear model adjusted R^2^= 0.94, p-value = 0.001 for model, post-vern p-value = 0.006, percentage survival to silique set p-value = 0.002, Fig. 7F, G).

## Discussion

Investigation of the dynamics of key floral integrators in the field has recently led to important insights into their molecular response to natural environments (Antoniou-Kourounioti et al., 2018; Song et al., 2018). Here we integrate experiments on the expression of *FLC* in field conditions with investigation on how natural genetic variation and induced mutation interact in distinct climates to affect phenotype and fitness. In our previous work, we found that, as expected from laboratory studies, *FLC* was repressed and *VIN3* was induced by winter cold (Antoniou-Kourounioti et al., 2018; Hepworth et al., 2018). However, in 2014-5, winter conditions in Norwich generated a subtly different response to those in Sweden, activating the VIN3-independent transcriptional shutdown before the VIN3-dependent epigenetic pathway. In the following season, this pattern was triggered again, not only in Norwich but also in North Sweden, implying that this is a common occurrence that plants must adjust to across climates, although with different frequencies at different locations.

We find that *FLC* level variation in both accessions and NILs is largely due to differences in the starting *FLC* level and the VIN3-independent rate of shutdown, which vary widely between accessions but not between climate sites or years (Fig. 5). This variation in response, as well as absolute levels of *FLC*, partly explains why *FLC* levels measured at any one time may not correlate with the final flowering time phenotype in different accessions (Sasaki et al., 2018).

In Norwich, we investigated the mechanisms of the responses. Contrary to our expectations, mutants of the autonomous pathway, despite upregulating *FLC* in *fri* plants in laboratory conditions (Ausín et al., 2004; Liu et al., 2010; Wu et al., 2016) generally behaved in a remarkably similar manner to Col *FRI* in terms of vernalization response (Fig. 2, 3). However, the reduced response of the *fve* mutants in the VIN3-independent phase, though not significant in each year, may be worth further investigation, as FVE has been implicated in intermittent cold-sensing through histone deacetylation at *FLC*, independently of vernalization (Kim et al., 2004; Jung et al., 2013).

All three factors that set *FLC* levels in the field (starting levels, VIN3-independent and VIN3-dependent phases) require chromatin modifiers for their correct function. Whether this is a direct effect on *FLC* chromatin, or due to epigenetic control of unidentified *trans* factors that govern the VIN3-independent response, is unclear. The VIN3-independent phase does not generate a strong epigenetic memory of cold, supporting a *trans* effect (Hepworth et al., 2018). However, the VAL1 transcription factor, which binds directly to *FLC* and is required for PRC2 action there (Qüesta et al., 2016; Yuan et al., 2016), has a strong effect on the VIN3-independent pathway (Fig. 3), suggesting it acts directly, whether via PRC2 or another mechanism.

These epigenetic factors are rarely found in genome-wide-association studies for flowering time, likely because their modification would have pleiotropic effects. In the natural accessions we find that the VIN3-dependent phase is indeed the least variable (Fig. 5). This may explain why so much variation in vernalization maps to *FLC* itself (Lempe et al., 2005; Sánchez-Bermejo et al., 2012; Dittmar et al., 2014; Sasaki et al., 2015; Bloomer and Dean, 2017; Sasaki et al., 2018).

The importance of *FLC* variation in adaptation for natural populations is well established (Méndez-Vigo et al., 2011; Sánchez-Bermejo et al., 2012; Dittmar et al., 2014; Li et al., 2014; Duncan et al., 2015; ågren et al., 2016; Bloomer and Dean, 2017). Nevertheless, we found that even in a challengingly warm year, across three climates vernalization requirement saturated well before midwinter (Fig. 6). This response had been seen previously in North Sweden in the locally adapted Löv-1 accession as a reaction to the extreme winters (Duncan et al., 2015). However, we find that this is a general response in all our tested accessions. The consequences of incomplete vernalization are severe – in our 2016-17 transfer experiments this led to delayed flowering and reduced fecundity (Fig. 6, S7-4). Hence, there is strong selection pressure on survival for RV *FLC* alleles in warm conditions, as demonstrated in field experiments in Italy (ågren et al., 2013; Grillo et al., 2013; Dittmar et al., 2014; ågren et al., 2016).

However, in reciprocal transplant experiments, ågren and coworkers found that the Italian RV *FLC* allele generally had a neutral effect on survival in Sweden, as expected for survival alleles when outside their adapted locality (Fournier-Level et al., 2011; ågren et al., 2013; ågren et al., 2016). Likewise, in 2014-5, the low mortality and synchronicity of flowering did not indicate any strong effect of the *FLC* allele after winter across our field sites (Fig. S6-2), suggesting that the autumn saturation of vernalization had removed its influence. Nevertheless, we observed that the Swedish, cold-winter-adapted accessions and SV alleles expressed the highest *FLC* levels during autumn (Fig. 2, 3, S2-1, S3-1). In North Sweden in 2016-7, an unusually warm growing season revealed an adaptive role for these high *FLC* levels. In plants with an SV allele, including the locally-adapted Swedish accessions, the higher level of *FLC* expression protected against precocious flowering and its consequent reduced survival (Fig. 7, S7-1). These rare, but highly selective occurrences, may be a driving force for local adaptation. That the effects of these alleles are only revealed occasionally is a logical consequence of the fact that the flowering time genes can have strong or weak effects depending on the environment (Wilczek et al., 2009; Burghardt et al., 2016; Fournier-Level et al., 2016; Taylor et al., 2019).

*FLC* also controls fecundity as well as survival (ågren et al., 2013). Li *et al*. (2014) found that SV *FLC* alleles produced lower seed weight compared to RV alleles in non-saturating vernalization conditions. In Arabidopsis, saturation of vernalization requirement is known to increase flowering branch production, particularly rosette branch production, and this effect is linked to *FLC* (Huang et al., 2013; Jong et al., 2019). In the Arabidopsis relative *Arabis alpina*, this effect has been linked to the *FLC* homologue *PEP1*, and has a subsequent effect on fitness by influencing silique production (Lazaro et al., 2018). We confirmed the effect of the *FLC* allele in *A. thaliana* on branching response to vernalization in laboratory conditions (Fig. S7-2). In the field, silique production, and hence fitness, in surviving plants was closely linked to branch production (Fig. 7). As expected, *FLC* levels in autumn did not correlate with branch production in spring (Fig. S7-1H), and in Norwich 2014-15, there was little variation in branch production. However, we found that under the conditions in the field in North Sweden, branch production was negatively related to a factor we named ‘*FLC* post-vern’, which encoded the relationship between *FLC* levels and subsequent flowering in warm controlled conditions (Fig. 7). Variation in this factor probably derives from regulatory differences at *FLC* post-cold, such as the reactivation phenotype of the Löv-1 *FLC* allele (Shindo et al., 2006; Coustham et al., 2012; Qüesta et al., 2020), allowing *FLC* repression of flowering to saturate at the shoot apical and axillary meristems at different rates, a phenomenon that occurs in the perennial relative *Arabis alpina* (Wang et al., 2009; Lazaro et al., 2018). Why the field conditions in Sweden, but not Norwich, revealed these conditions is not yet clear. This property of *FLC* appears to be regulated separately to that of *FLC* vernalization response, as a slow VIN3-independent shutdown does not necessarily cause slow flowering in the warm. As such, it is an example of how one gene can directly regulate independent phenotypes in response to different environmental cues, mitigating evolutionary constraints in which selection for one phenotype may be traded-off against concomitant changes to another (Auge et al., 2019). Given that in many cases there is stronger selection for high branch production than flowering time in the field (Taylor et al., 2019), this is likely to be an important evolutionary constraint for *FLC* alleles.

In summary, our detailed analysis of the different phases of *FLC* silencing through winters in distinct climates, over multiple years, has given a clear picture of the mechanistic basis of adaptation in vernalization response. *FLC* starting levels and early phases of *FLC* silencing are the major determinants for variation in vernalization. Non-coding *FLC* SNP variation aligns vernalization response to different climatic conditions and year-on-year fluctuations in natural temperature profiles. In a changing climate, understanding the complex genotype by environment interactions that govern timing mechanisms will become ever more important.

## Materials and Methods

### Plant materials

Sources of previously described mutant lines and transgenics are presented in Supplementary Table 1.

### NILs

All near-isogenic lines were produced by six rounds of backcrossing to the Col *FRI* parent, selecting for the introgressed *FLC* in each generation, before one round of selfing and selection of homozygous families.

**Supplementary Table 1.** Sources of previously published mutants and transgenics.

| Genotype | Details | Norwich 2014 | South Sweden 2014 | North Sweden 2014 | Norwich 2016 | South Sweden 2016 | North Sweden 2016 | Source |
| --- | --- | --- | --- | --- | --- | --- | --- | --- |
| Col *FRI* | Col-0 with *FRI* from San Feliu introgressed | X | X | X | X | X | X | Lee and Amasino (1995) |
| **Accessions** |  |  |  |  |  |  |  |  |
| Col-0 |  | X | X | X | X | X | X |  |
| Bro1-6 |  | X | X | X | X | X | X |  |
| Edi-0 |  | X | X | X | X | X | X |  |
| Löv-1 |  | X | X | X | X | X | X |  |
| Ull2-5 |  | X | X | X | X | X | X |  |
| Var2-6 |  | X | X | X | X | X | X |  |
| **Near Isogenic Lines** |  |  |  |  |  |  |  |  |
| Bro NIL | *FLC* from Bro1-6 backcrossed to Col *FRI* background six times. |  |  |  | X |  |  | This paper |
| Edi NIL |  | X | X | X | X | X | X | This paper |
| Löv NIL1 |  | X | X | X | X | X | X | Duncan et al. (2015) |
| Löv NIL2 |  | X | X | X | X | X | X | Duncan et al. (2015) |
| Ull NIL | *FLC* from Ull2-5 backcrossed to Col *FRI* background six times. |  |  |  | X | X | X | This paper, derived from Strange et al. (2011) |
| Var NIL |  | X |  |  | X | X | X | Li et al. (2015) |
| **Mutants** |  |  |  |  |  |  |  |  |
| *vin3-1 FRI* |  |  |  |  | X | X | X | Sung & Amasino (2004) |
| *vin3-4 FRI* |  | X |  |  | X |  |  | Bond et al. (2009) |
| *vrn1-4 FRI* |  | X |  |  |  |  |  | Sung & Amasino (2004) |
| *vrn2-1 FRI* |  | X |  |  | X |  |  | Yang et al. (2017) |
| *vrn5-8 FRI* |  |  |  |  | X |  |  | Greb et al. (2007) |
| *ndx1-1 FRI* |  | X |  |  | X |  |  | Sun et al. (2013) |
| *fca-9* |  | X |  |  |  |  |  | Liu et al. (2007) |
| *fld-4* |  | X |  |  |  |  |  | Liu et al. (2007) |
| *fve-3* |  | X |  |  |  |  |  | Ausín et al. (2004) |
| *fca-9 FRI* | Cross Col *FRI* and lines reported above |  |  |  | X |  |  | This paper |
| *fld-4 FRI* | Cross Col *FRI* and lines reported above |  |  |  | X |  |  | This paper |
| *fve-3 FRI* | Cross Col *FRI* and lines reported above |  |  |  | X |  |  | This paper |
| *val1-2 FRI* |  |  |  |  | X |  |  | Qüesta et al. (2016) |
| *sdg8 FRI* |  |  |  |  | X |  |  | Yang et al. (2014) |
| *lhp1-3 FRI* |  |  |  |  | X |  |  | Mylne et al. (2006) |

### Experimental conditions

#### Field experiments

Field experiments have been described previously (Antoniou-Kourounioti et al., 2018 (2016-7 winter); Hepworth et al., 2018 (2014-5 winter)). Briefly, seeds were stratified at 4°C for three days. For gene expression measurements, for all field sites and sowing dates and timepoints within them, six replicate tray-cells were sown using a block-randomised design within 5.7cm 28-cell trays (Pöppelman, Lohne, Germany), and where each replicate included material from at least three plants. For flowering time, plants were thinned to a single plant per cell in 3.9cm 66-cell trays (Pöppelman). Trays were watered when necessary.

In Norwich, trays were placed on benches in an unheated, unlit glasshouse, and bedded in vermiculite. For expression, plants were randomised within 6 single-replicate sample-sets, which were then randomised using Research Randomiser (Urbaniak and Plous, 2015) in a 3 complete block design lengthwise along the greenhouse, adjusting to ensure each of the two replicates per block were on different benches. For flowering time, plants were block-randomised per tray and per block.

In Sweden, trays were germinated and grown outside under plastic covers for two weeks at Mid Sweden University, Sundsvall (North Sweden) or Lund University (South Sweden). Trays were then moved to the experiment sites dug into the soil. The experiment site in the North was at Ramsta (62° 50.988’N, 18° 11.570’E), and in the South at Ullstorp (56° 06.6721’N, 13° 94.4655’E). Expression sample-sets were randomised in three blocks. Flowering time plants were in a completely randomised design across two (2014-15) or three (2016-17) blocks.

Plants were sown and moved to the field site on: Norwich first year, sown into position on 29^th^ September 2014, second year, sown into position on 15th September 2016: South Sweden first year, sown 24^th^ September 2014, moved 8^th^ October 2014, second year sown on 6th September 2016, moved on 21st September 2016: North Sweden first year, first planting ‘A’ sown 26^th^ August 2014, moved 11^th^ September 2014; second planting ‘B’ sown 8^th^ September 2014, moved 24^th^ September 2014: North Sweden second year, sown on 12th August 2016, and moved 24th August 2016.

For the transferred plants, in Norwich 2016-7, for each transfer 6 trays (each holding 2 replicates, total n=12) were moved from the unheated, unlit, ventilated greenhouse to a greenhouse with supplementary lighting (600W HPS lamps) and heating set to 22°C/18°C, 16 light/8 hour dark, and 70% humidity on 21^st^ October 2016, 3^rd^ November 2016, 17^th^ November 2016, 30^th^ November 2016, 21^st^ December 2016 and 26^th^ January 2017. For selected time points, plants were covered with ventilated clear plastic bags to collect seed for weighing. In South Sweden, trays were transferred from the field site to heated, lit greenhouses at Lund University, on 1^st^ October, 22^nd^ October, 19^th^ November and 17^th^ December 2016. In North Sweden, for each transfer 3 trays (each holding 4 replicates, n=12) were moved from the field site to a greenhouse set to 16 hr light, 22°C, at Midsweden University, Sundsvall (as in Duncan et al., 2015), on 6^th^ September, 4^th^ October, 1^st^ November and 24^th^ November 2016.

For all expression analysis except the 48hr sampling (Fig. S3-3), 6 replicates per timepoint per genotype were chosen in order to allow sufficient number of samples for statistical analysis while allowing for losses in the field and to allow duplication within randomization blocks. Where resulting samples are smaller, this is due to experimental or processing loss (e.g., death of plants in the field, degradation due to poor sample quality or processing, see RNA extraction and QPCR). For Fig. S3-3, 3 replicates per timepoint per genotype were chosen due to space constraints. Each expression sample (single replicate) was of at least 3 plants pooled. For flowering time, 12 plants per genotype per transfer condition were chosen to provide replication across blocks and trays while remaining within size constraints. For field flowering in North Sweden 2016-17, 36 plants per genotype were sown to allow for losses, although in the event these were more substantial than anticipated.

Temperature was recorded at plant level at each site with TinyTag Plus 2 dataloggers (Gemini Data Loggers (UK) Ltd, Chichester, UK). Bolting was scored when flower buds were visible at the shoot apical meristem. For the North Sweden 2016-7 field experiment, plants were scored for survival and flowering in the field from planting to December 2016, and then from March to May 2017. Plants were harvested and scored for branching and silique production after the end of flowering, in July 2017.

#### Branching analysis

Fig. S7-2, S7-3; Seeds were sown on soil, stratified after sowing for three days at ∼4-5°C, and transferred to a Norwich long-day glasshouse set to 18°C/15°C, 16/8 hour light/dark conditions for 7 days before being returned to vernalization conditions (a 4°C growth chamber under short day, low light conditions; 8/16 hour light/dark) for 12, 8, 4, and 0 weeks. Sowing was staggered so that after vernalization all plants were transferred to glasshouse conditions simultaneously. Plants were scored for their flowering time, total branch number, cauline branch number and rosette branch number. In all cases, plants were randomized into blocks and at least three replicate plants for each accession/cultivar per treatment were scored for their flowering and branching phenotypes. Primary rosette and cauline branch number were scored at senescence.

### RNA extraction and QPCR

RNA extraction and QPCR for field experiments were performed as described in Hepworth *et al*. (2018) and Antoniou-Kourounioti *et al*. (2018). Field data, was unified across sites and timepoints within the yearly datasets and normalised to a synthetic control sample, as described in Hepworth et al. (2018). QPCR results were analysed using LinReg (Ruijter et al., 2009), and normalised to the geometric means of At5g25760 (‘*PP2A*’) and At1g13320 (‘*UBC’*) control genes (Czechowski et al., 2005; Yang et al., 2014).

QPCR samples that showed high Cp values (UBC Cp>28 for LinReg analysis) of the control genes, indicating possible degradation, were excluded if test amplicons (FLC, VIN3) results were also anomalous (criteria: completely absent, or varying from non-flagged samples by an order of magnitude). Any measurements where amplicon Ct values varied by more than 0.6 were also excluded if insufficient sample was available for a repeat.

Primers used are described in Supplementary Table 2.

**Supplementary Table 2.**
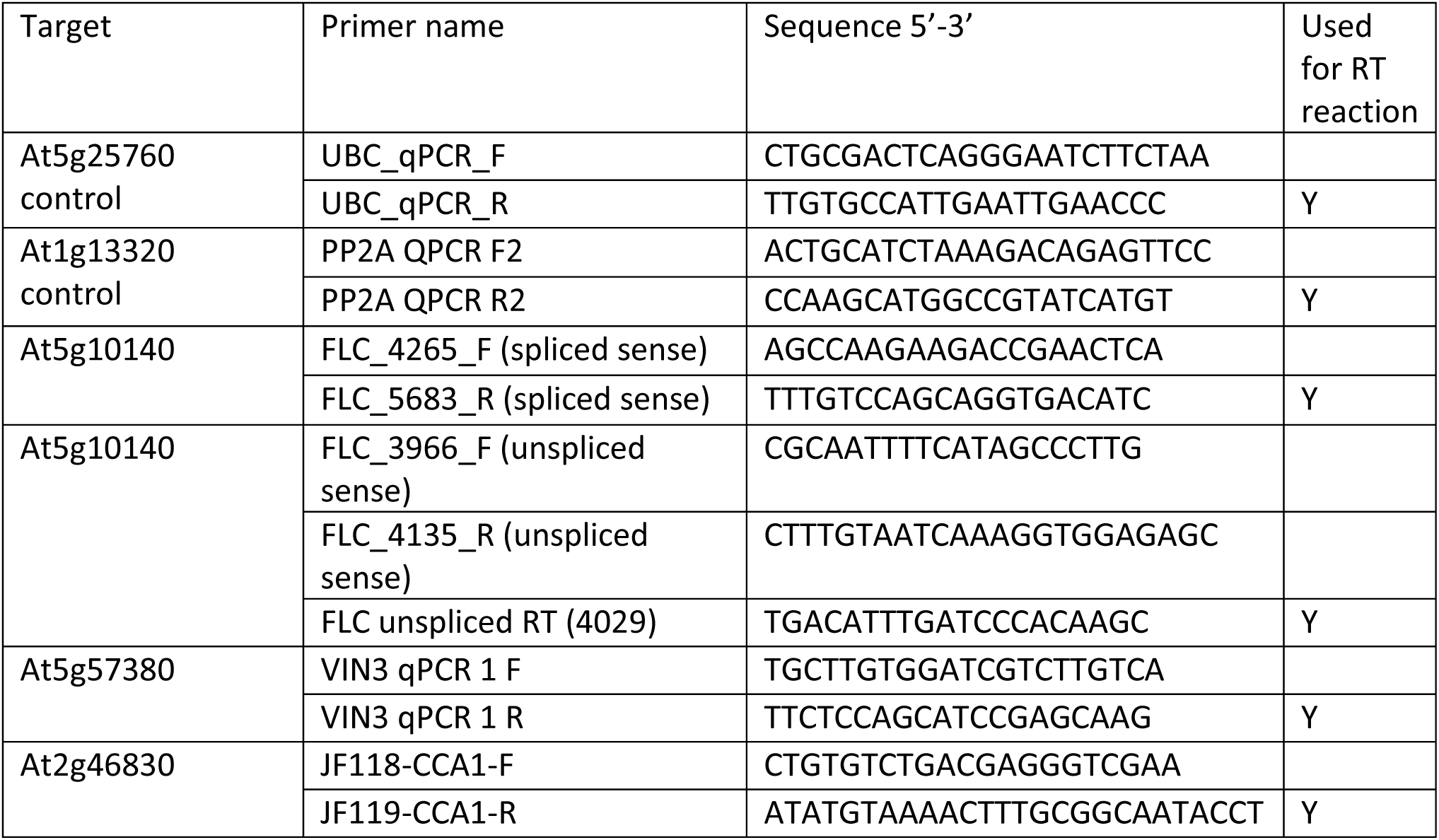
Primers used for PCR.

### Statistics

For the analysis presented in Supplementary file 1 and Figures 2-4, the Starting level comparisons and the Slope comparisons were done separately, using the following methods. For the starting levels, ANOVA was performed in the R (R Core Team, 2018) statistical language using the lm function, followed by a Dunnett post hoc test comparing all genotypes against the control Col *FRI*. For the slopes, the lmer function from the lmerTest package in R was used to perform the same comparison on the genotype-timepoint interaction, controlling for blocks as random factors.

We used the R package cvequality (Version 0.1.3; Marwick and Krishnamoorthy, 2019) with the asymptotic test (Feltz and Miller, 1996) to assess differences between the coefficients of variation of different groups as described in the text and Supplementary Table 3. The significance limit was adjusted to control the false discovery rate using the Benjamini-Hochberg procedure with a false discovery rate of 0.05 (Supplementary Table 3, Supplementary file “coef_var_comparison_analyses.xlsx”). This analysis was performed including all the natural accession and NILs combined (Fig. 4), but also for all the accessions separately and for all the NILs separately (including Col *FRI* in both cases). The different analyses do not change our overall conclusions and all three are reported in Supplementary Table 3.

For multiple regression on field data, R was used to obtain minimal adequate models using linear regression (lm function), except when n>10 for count data, for which general linear models (GLM, glm function) using Poisson error distributions were used (total numbers of siliques), or for proportion data for which GLMs with binomial errors were used (survival, bolting before winter).

**Supplementary Table 3.** List of all comparisons of coefficients of variation. Differences between the coefficients of variation of different groups (sites, features, years; 24 comparisons).

| # | Comparison | Groups | Signif. Acc | Signif. NILs | Signif. combined |
| --- | --- | --- | --- | --- | --- |
| 1 | All | VIN3-independent in North Sweden 2016-7,<br>VIN3-independent in Norwich 2014-5,<br>VIN3-independent in Norwich 2016-7,<br>Starting levels in North Sweden 2016-7,<br>Starting levels in Norwich 2014-5,<br>Starting levels in Norwich 2016-7,<br>Starting levels in North Sweden 2014-5 1st planting,<br>Starting levels in North Sweden 2014-5 2nd planting,<br>Starting levels in South Sweden 2014-5,<br>VIN3-dependent in North Sweden 2016-7,<br>VIN3-dependent in Norwich 2014-5,<br>VIN3-dependent in Norwich 2016-7,<br>Combined rate in North Sweden 2014-5 1st planting, | * | * | * |
|  |  | Combined rate in North Sweden 2014-5<br>2nd planting,<br>Combined rate in South Sweden 2014-5 |  |  |  |
| 2 | Sites (features<br>combined) | North Sweden 2016-7, Norwich 2014-5,<br>Norwich 2016-7, North Sweden 2014-5 1st<br>planting, North Sweden 2014-5 2nd<br>planting, South Sweden 2014-5 |  |  |  |
| 3 | Features (2014<br>sites combined) | VIN3-independent, Starting levels, VIN3-dependent, Combined rate | * | * | * |
| 4 | Features (2016<br>sites combined) | VIN3-independent, Starting levels, VIN3-dependent | * | * | * |
| 5 | Features (sites<br>combined) | VIN3-independent, Starting levels, VIN3-dependent, Combined rate | * | * | * |
| 6 | Features in<br>Norwich 2014-5 | VIN3-independent, Starting levels, VIN3-dependent | * | * | * |
| 7 | Features in<br>Norwich 2016-7 | VIN3-independent, Starting levels, VIN3-dependent |  | * | * |
| 8 | Features in<br>Norwich | VIN3-independent, Starting levels, VIN3-dependent | * | * | * |
| 9 | VIN3-independent<br>and Starting<br>levels in<br>Norwich | VIN3-independent, Starting levels |  | * | * |
| 10 | VIN3-independent<br>and VIN3-dependent in<br>Norwich | VIN3-independent, VIN3-dependent | * | * | * |
| 11 | VIN3-dependent<br>and Starting<br>levels in<br>Norwich | Starting levels, VIN3-dependent |  |  | * |
| 12 | Features in<br>North Sweden<br>2014-5 1st<br>planting | Starting levels, Combined rate | * | * | * |
| 13 | Features in<br>North Sweden<br>2014-5 2nd<br>planting | Starting levels, Combined rate | * |  | * |
| 14 | Features in<br>South Sweden<br>2014-5 | Starting levels, Combined rate | * |  | * |
| 15 | Features in<br>North Sweden<br>2016-7 | VIN3-independent, Starting levels, VIN3-dependent |  |  | * |
| 16 | Features in<br>Sweden 2014-5 | Starting levels, Combined rate | * | * | * |
| 17 | Features in<br>Sweden | VIN3-independent, Starting levels, VIN3-dependent, Combined rate | * | * | * |
| 18 | Starting levels | Starting levels in North Sweden 2016-7,<br>Starting levels in Norwich 2014-5,<br>Starting levels in Norwich 2016-7,<br>Starting levels in North Sweden 2014-5 1st<br>planting,<br>Starting levels in North Sweden 2014-5 2nd<br>planting,<br>Starting levels in South Sweden 2014-5 |  |  |  |
| 19 | Shutdown rates | VIN3-independent in North Sweden 2016-7,<br>VIN3-independent in Norwich 2014-5,<br>VIN3-independent in Norwich 2016-7,<br>VIN3-dependent in North Sweden 2016-7,<br>VIN3-dependent in Norwich 2014-5,<br>VIN3-dependent in Norwich 2016-7,<br>Combined rate in North Sweden 2014-5 1st<br>planting,<br>Combined rate in North Sweden 2014-5<br>2nd planting,<br>Combined rate in South Sweden 2014-5 |  | * | * |
| 20 | Combined<br>shutdown rates<br>(Sweden 2014-5) | Combined rate in North Sweden 2014-5 1st<br>planting,<br>Combined rate in North Sweden 2014-5<br>2nd planting,<br>Combined rate in South Sweden 2014-5 |  |  |  |
| 21 | VIN3-independent<br>rates | VIN3-independent in North Sweden 2016-7,<br>VIN3-independent in Norwich 2014-5,<br>VIN3-independent in Norwich 2016-7 |  |  |  |
| 22 | VIN3-dependent<br>rates | VIN3-dependent in North Sweden 2016-7,<br>VIN3-dependent in Norwich 2014-5,<br>VIN3-dependent in Norwich 2016-7 |  |  |  |
| 23 | Shutdown rates<br>(sites combined) | VIN3-independent, VIN3-dependent,<br>Combined rate | * | * | * |
| 24 | VIN3-independent<br>and Starting<br>levels (sites<br>combined) | VIN3-independent, Starting levels | * | * | * |

## Acknowledgements

For genetic materials, we are indebted to Prof. Johanna Schmitt (University of California Davis) for the kind gift of *vin3-1 FRI*, and Huamei Wang for creating the Ull and Bro NIL lines and preparing seed stocks. For the Swedish field sites, many thanks to family Öhman and Nils Jönsson. Thanks to Prof. James Brown (JIC) for advice on the statistics, and to Ian Baldwin (Senior Editor at eLIFE) for helpful suggestions on the initial submission of the manuscript. Finally the authors would like to reiterate their appreciation of all those from the Dean, Howard, Holm, Säll and Irwin groups who helped in cold, heat, wind, rain and laboratory with the field studies.

## Data Sources

Supplementary File 1 – Statistics for Fig. 2, 3 and 4.

Source Data 1 – Field temperatures

Source Data 2 – RNA Expression for all field experiments 2014-15

Source Data 3 – RNA Expression for all field experiments 2016-17

Source Data 4 – Flowering time and phenotypes for all field experiments 2014-15

Source Data 5 – Flowering time and phenotypes for all transfer experiments 2016-17

Source Data 6 – Flowering time. survival and phenotypes for North Sweden field experiments 2016-17

Source Data 7 – Flowering time and branching for accessions and NILs in constant-condition vernalisation treatments.

## Supplementary Information

**Figure S2-1.**
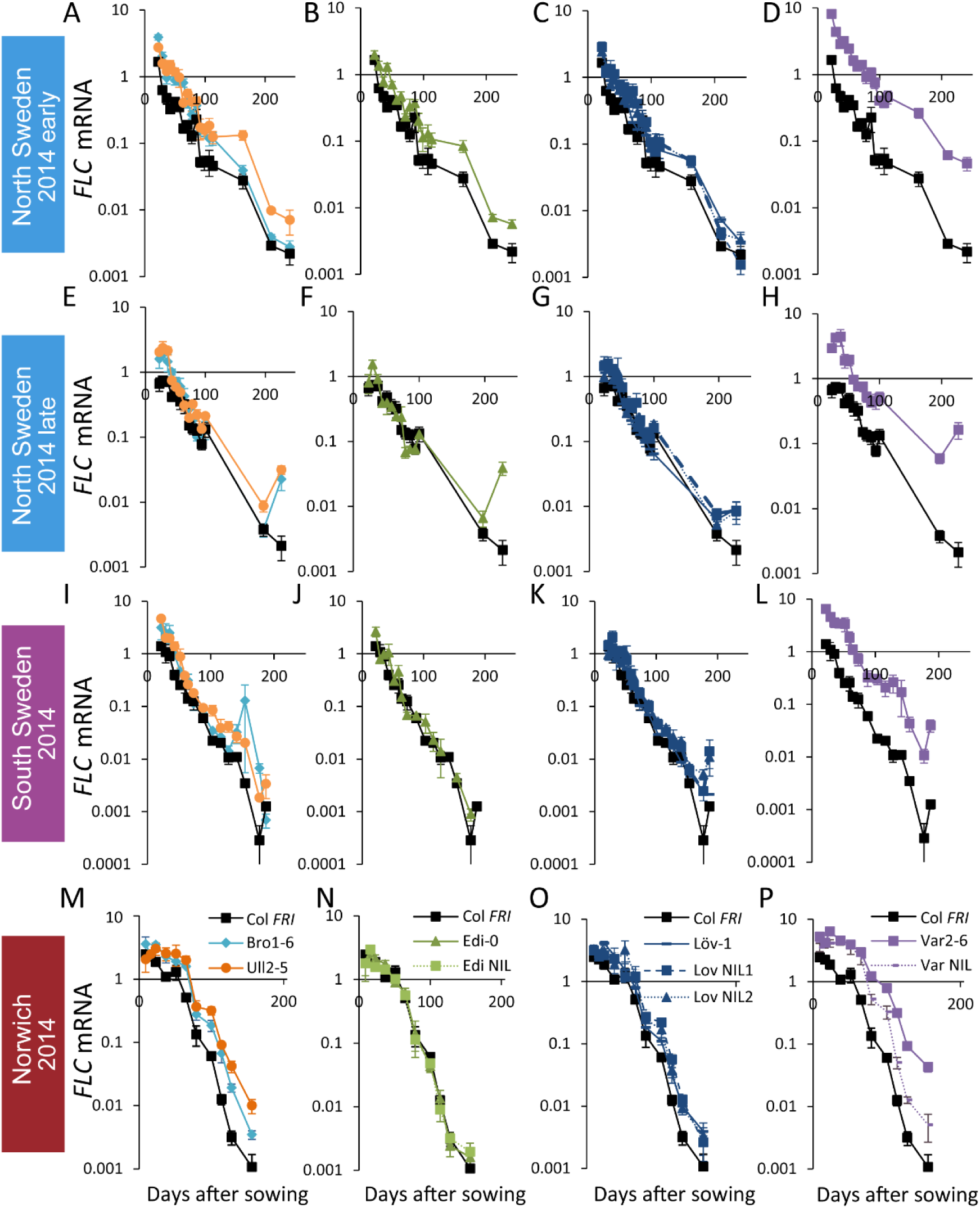
*FLC* downregulation in accessions and NILs in Norwich, North Sweden and South Sweden 2014-5. Expression normalised to control sample for 2014-5 (see Methods). (A-D) Norwich, (E-H) South Sweden, (I-L) North Sweden first planting, (M-P) North Sweden second planting. N=6 except where samples were lost to death or degradation (see Methods and Source Data 2). Error bars show s.e.m.

**Figure S2-2.**
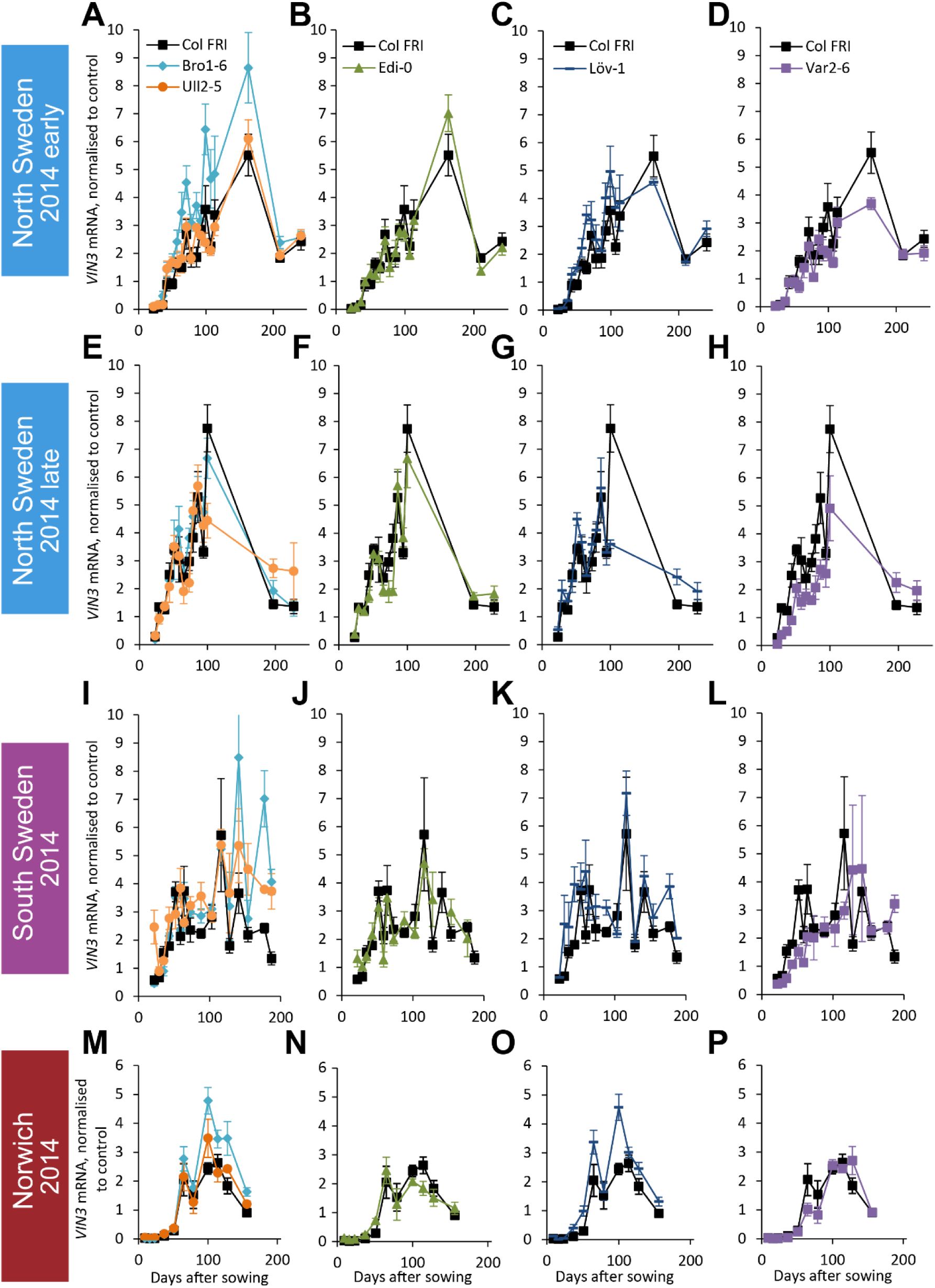
*VIN3* upregulation in accessions in Norwich, North Sweden and South Sweden 2014-5. *VIN3* expression normalised to control sample for 2014-5 (see Methods). N=6 except where samples were lost to death or degradation (see Methods and Source Data 2). Error bars show standard error of the mean (s.e.m).

**Figure S2-3.**
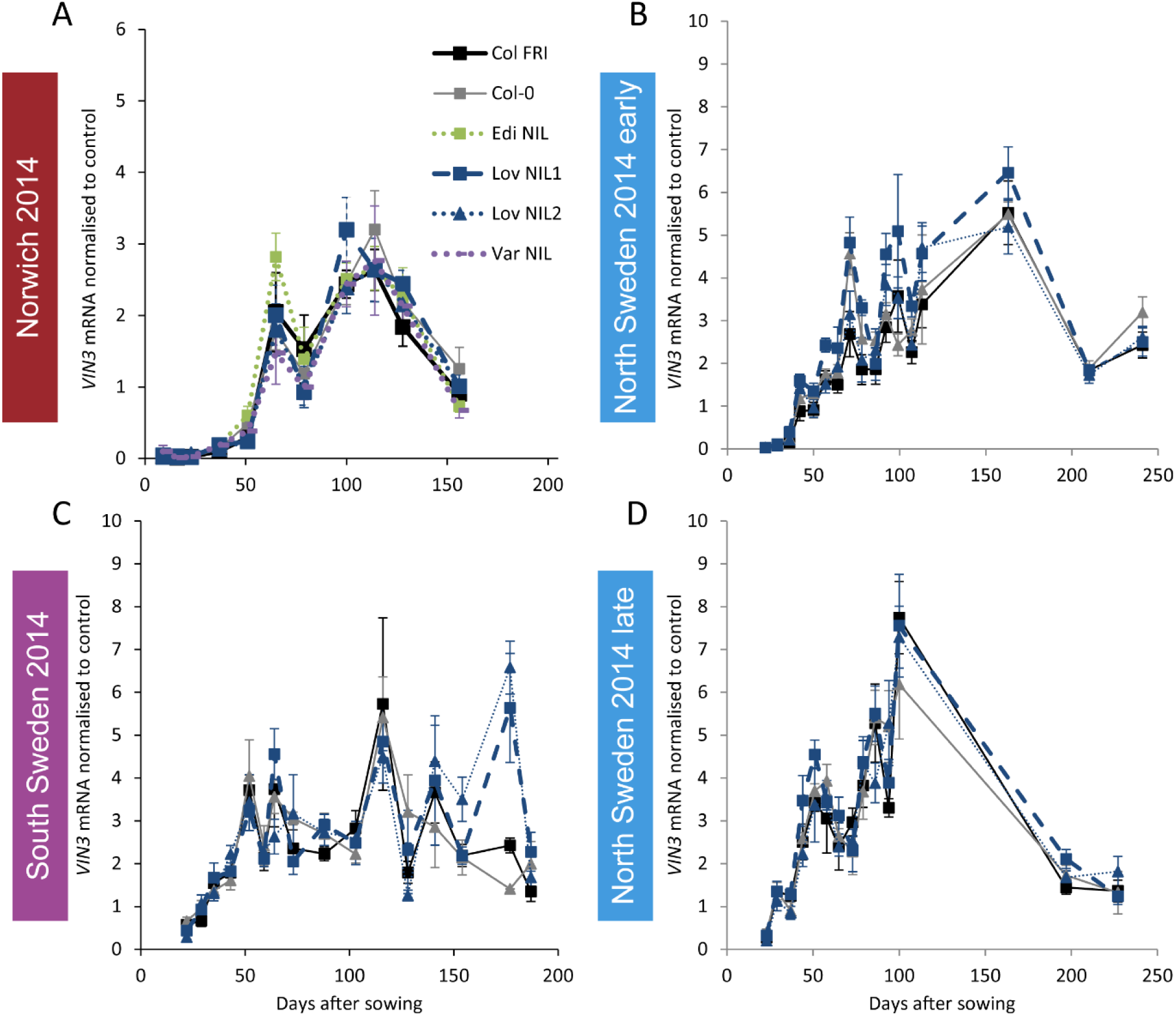
Expression of *VIN3* in NILs with the Col-0 *VIN3* allele in the field in 2014-2015. (A) Norwich, (B) North Sweden first planting, (C) South Sweden, (D) North Sweden second planting. N=6 except where samples were lost to death or degradation (see Methods and Source Data 2). Error bars show s.e.m.

**Figure S3-1.**
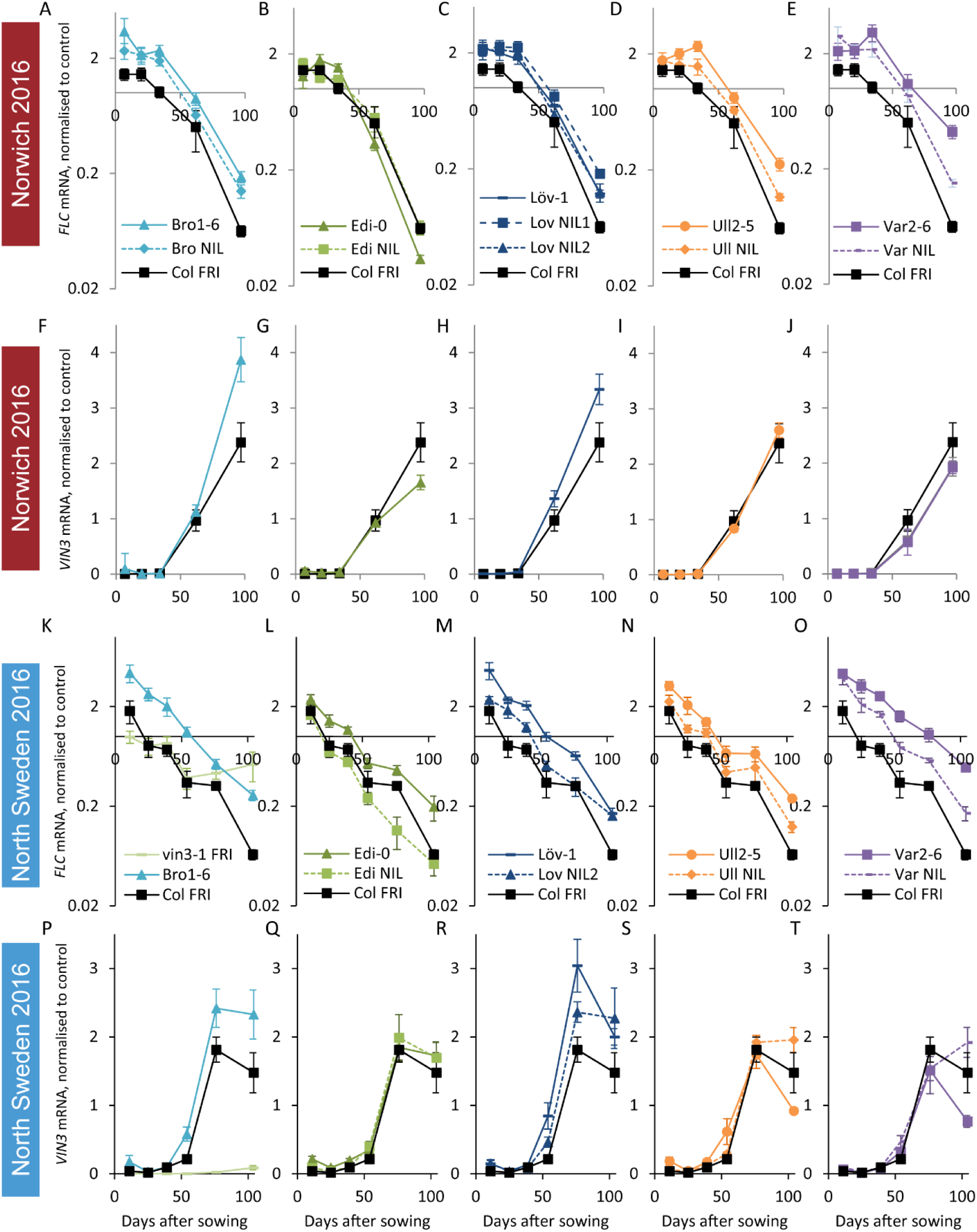
*FLC* downregulation and *VIN3* upregulation in accessions in Norwich and North Sweden in autumn/winter 2016. Expression normalised to control sample for 2016-7 (see Methods). (A-E) Norwich *FLC* mRNA, (F-J) Norwich *VIN3* mRNA, (K-O) North Sweden *FLC* mRNA, (P-T) North Sweden *VIN3* mRNA. N=6 except where samples were lost to death or degradation (see Methods and Source Data 3). Error bars show s.e.m.

**Figure S3-2.**
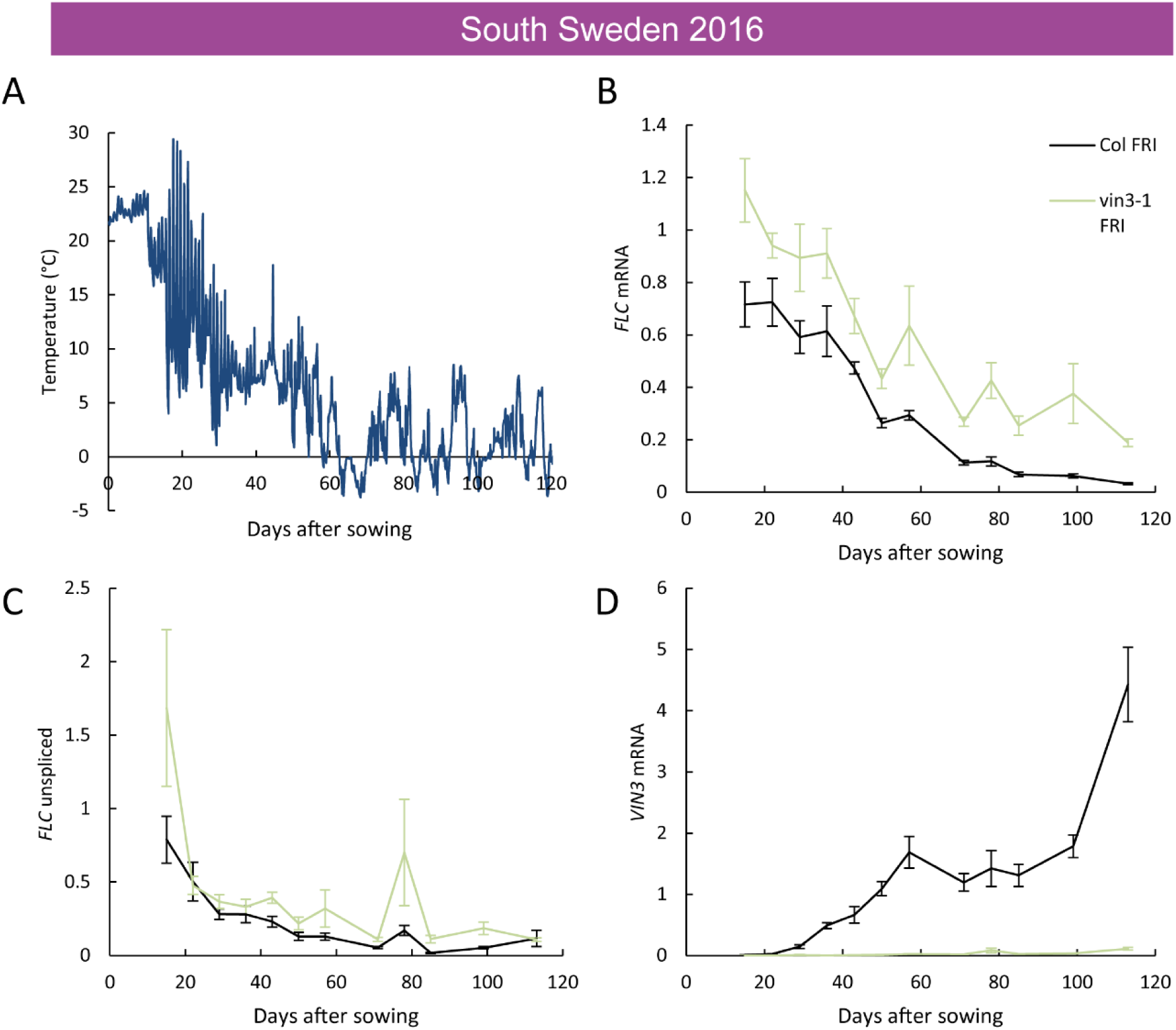
Downregulation of *FLC* and upregulation of *VIN3* in South Sweden in 2016. (A) Hourly temperature readings from plant-level in South Sweden 2016. (B) *FLC* mRNA levels in the Col *FRI* and *vin3-1 FRI* accessions over autumn, with *vin3-1* showing less repression, especially later in the season. (C) unspliced *FLC* levels. (D) *VIN3* mRNA levels, with *vin3-1 FRI* showing no induction. Expression normalised to control sample for 2016-7 (see Methods). N=6 except where samples were lost to death or degradation (see Methods and Source Data 3). Error bars show s.e.m.

**Figure S3-3.**
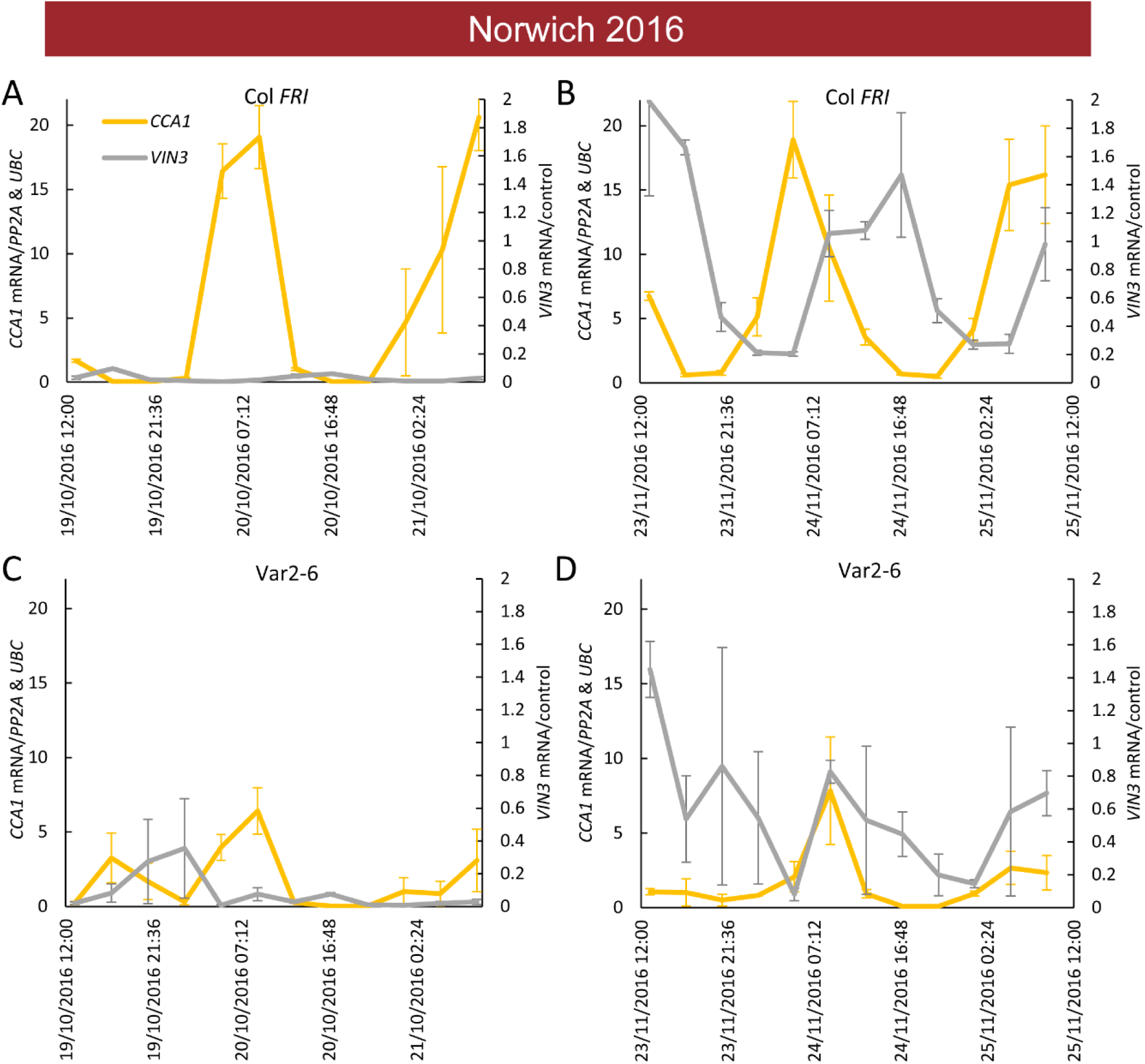
Low *VIN3* upregulation in Var2-6 is correlated with perturbation of the circadian clock. *VIN3* and *CCA1* expression measured over 48 hours in the field glasshouse in Norwich in 2016 in Col *FRI* and the Var2-6 accession. *CCA1* shows a circadian pattern throughout autumn in Col *FRI*, as does *VIN3* when it is upregulated later in the year. In Var2-6, *CCA1* expression is low, as is *VIN3* expression later in the year. (A) Expression in Col *FRI* in October. (B) Expression in Col *FRI* in November. (C) Expression in Var2-6 in October. (D) Expression in Var2-6 in November. N=3, Source Data 3. Error bars show s.e.m.

**Figure S4-1.**
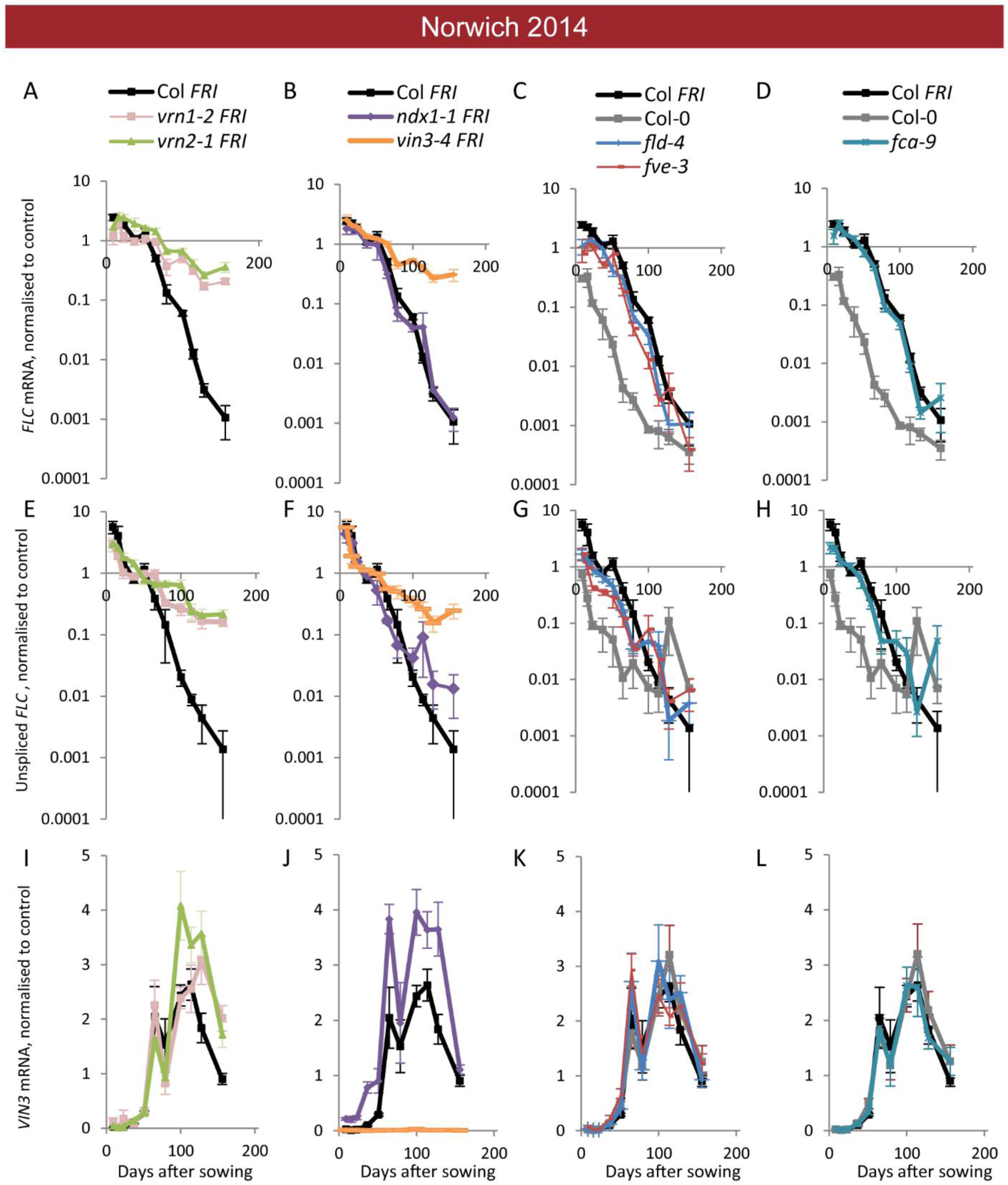
Expression of *FLC* and *VIN3* in all mutants in the field in Norwich 2014-2015. Expression normalised to control sample for 2014-5 (see Methods). (A-D) *FLC* mRNA, (E-H) unspliced *FLC* transcript, (I-L) *VIN3* mRNA. N=6 except where samples were lost to death or degradation (see Methods and Source Data 2). Error bars show s.e.m.

**Figure S4-2.**
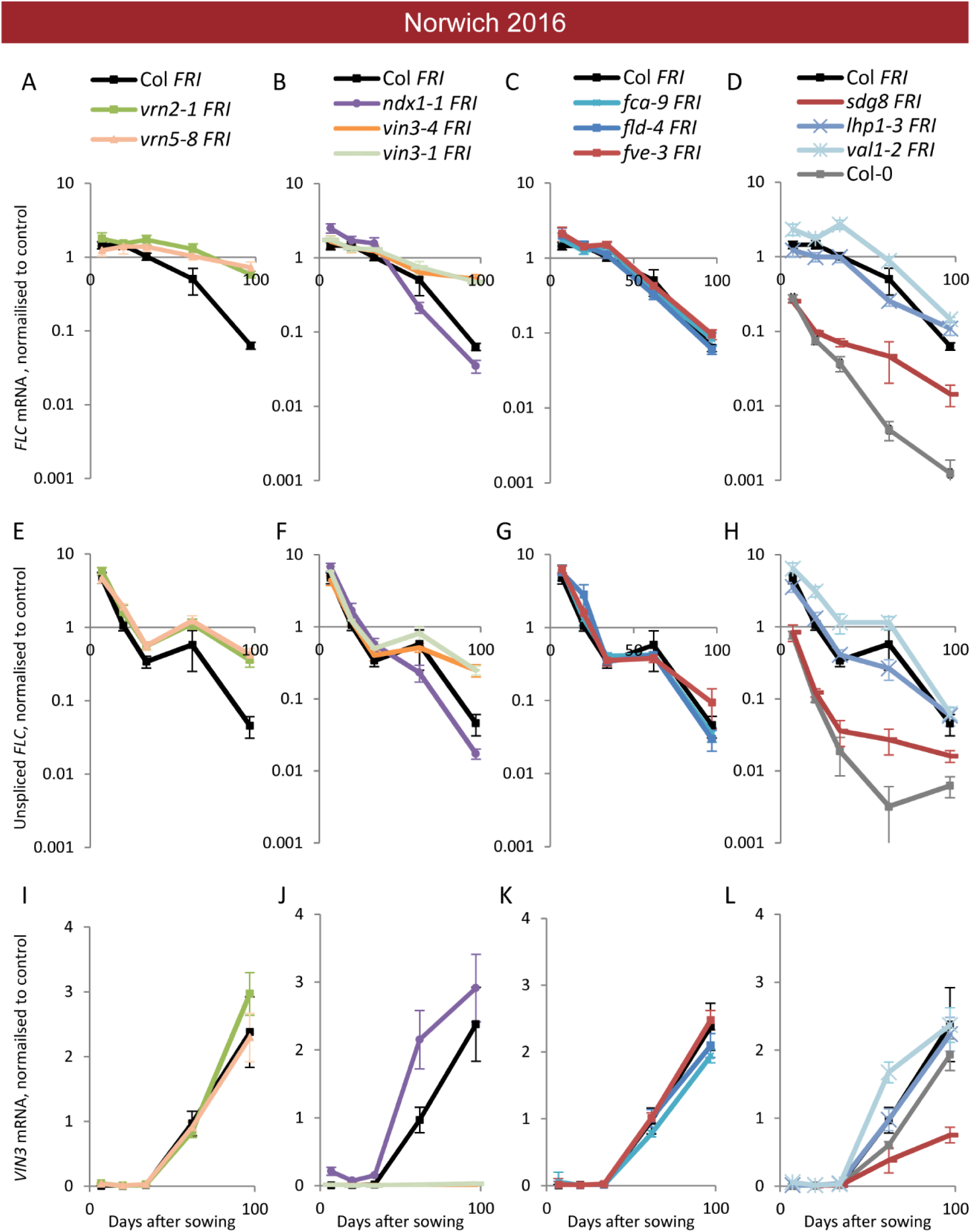
Expression of *FLC* and *VIN3* in all mutants in the field in Norwich 2016-2017. Expression normalised to control sample for 2016-7 (see Methods). (A-D) *FLC* mRNA, (E-H) unspliced *FLC* transcript, (I-L) *VIN3* mRNA. N=6 except where samples were lost to death or degradation (see Methods and Source Data 2). Error bars show s.e.m.

**Figure S6-1.**
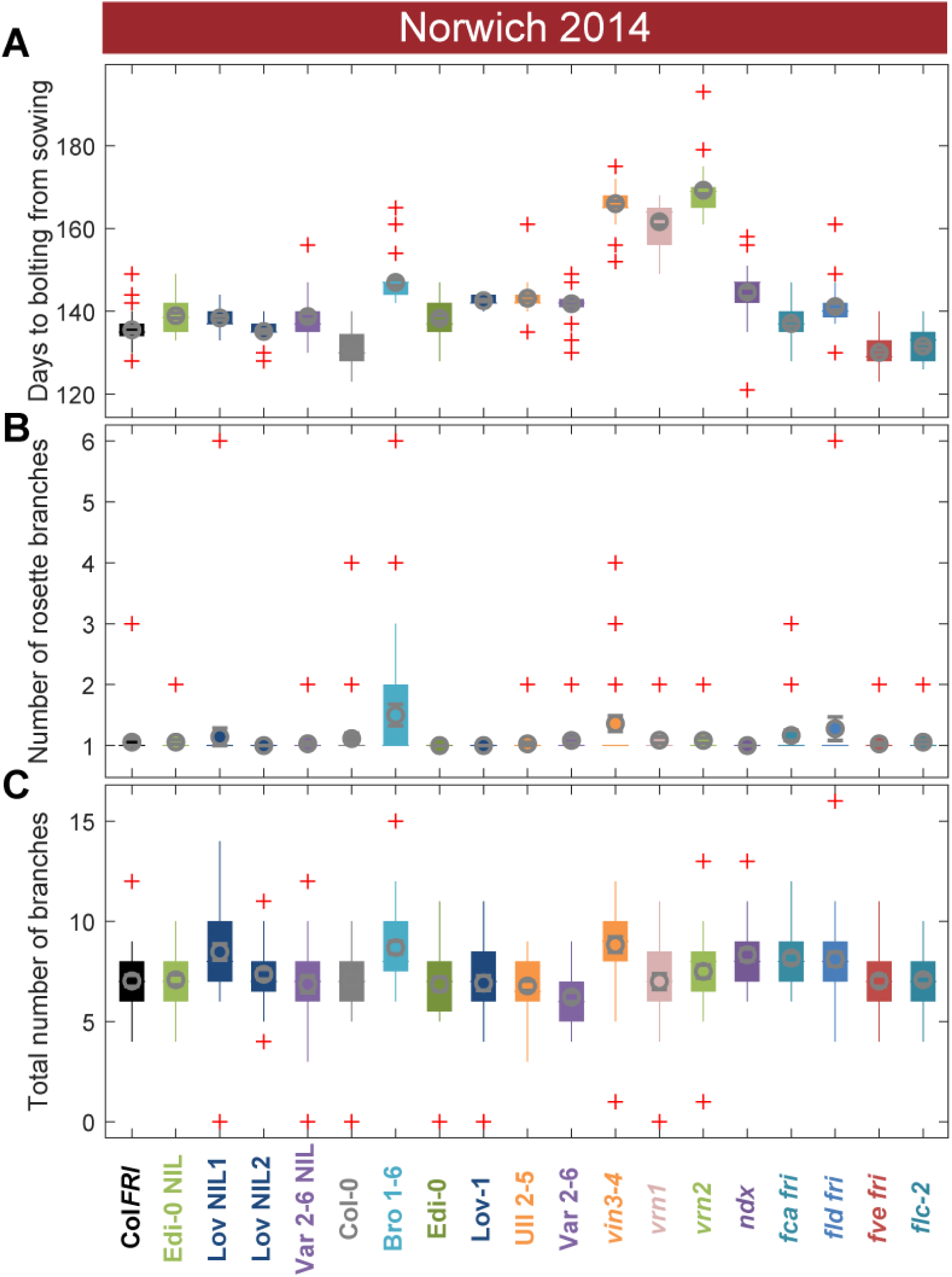
Flowering after winter in Norwich 2014-5 in the field was largely synchronous. **(A)** Time to bolting for each genotype in the ‘field’ glasshouse in Norwich 2014-5 experiment. **(B)** Number of rosette branches for plants shown in A. **(C)** Number of rosette and cauline branches for plants shown in A. Box-and-whiskers plot for time to bolting for each genotype, showing mean (grey circle), interquartile range (box), range (whiskers) and outliers (red crosses, values more than 1.5 times the interquartile range outside of the interquartile range). N=12, except where plants died, Source Data 4. Survival data for Norwich 2014-2015 not shown as only 2 plants died.

**Figure S6-2.**
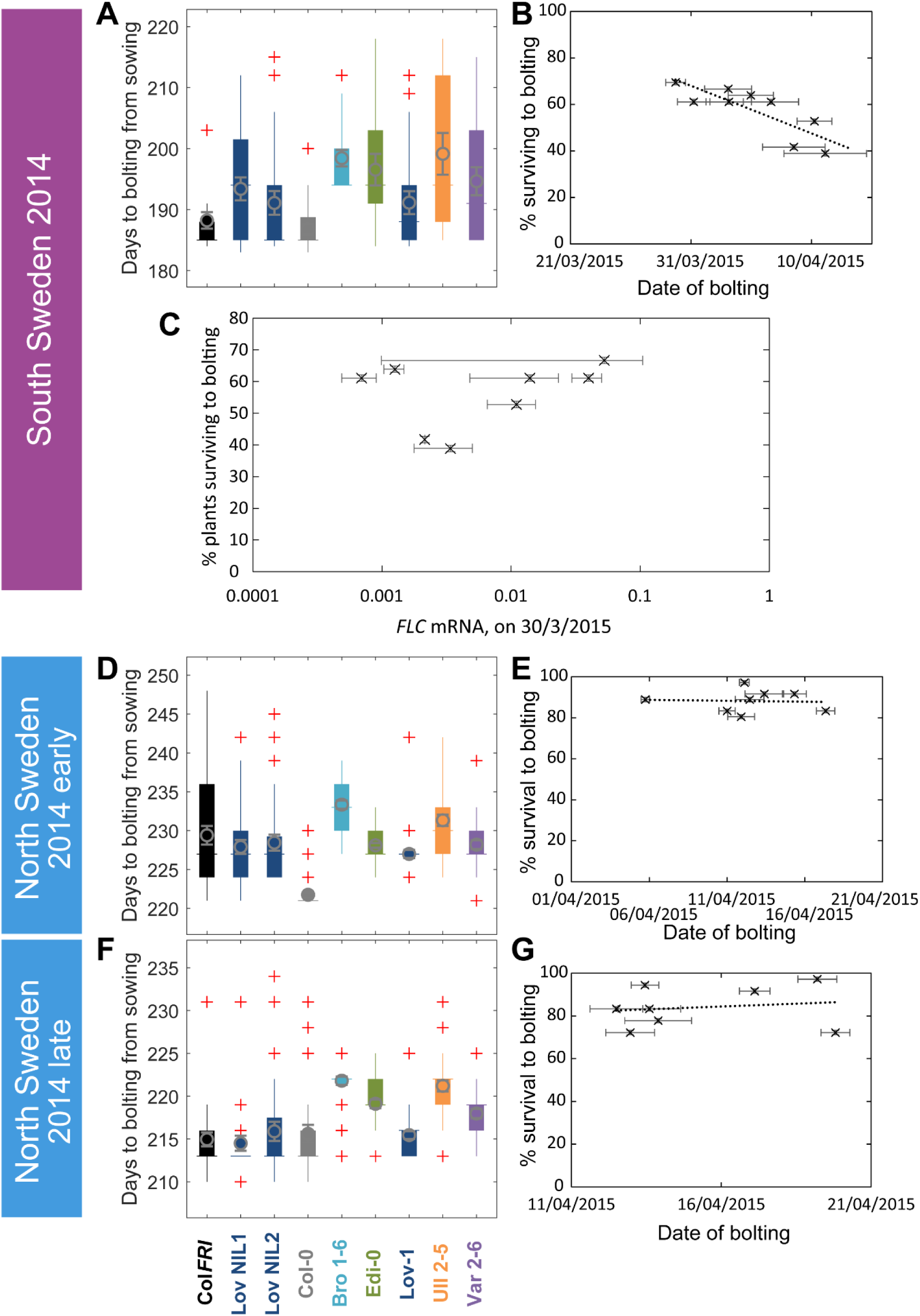
The transition to flowering after natural winters in South and North Sweden 2014-5 in the field was largely synchronous, while later bolting had a negative effect on survival only in South Sweden. **(A, D, F)** Box-and-whiskers plot for time to bolting for each genotype, showing mean (grey circle), interquartile range (box), range (whiskers) and outliers (red crosses, values more than 1.5 times the interquartile range outside of the interquartile range). **(B, E, G)** Percentage of germinated plants of each genotype surviving to date of bolting, plotted against the mean date of bolting for that genotype. **(B)** South Sweden, Generalised Linear Models (GLM) for binomial distribution, survival versus date of bolting, p=0.0416. **(C)** South Sweden, percentage survival versus mean *FLC* mRNA per genotype (normalised to control sample for 2014-5) on 30^th^ March, GLM for binomial distribution, ns. **(E, G)** GLM for binomial distribution, ns. N=12, except where plants died, Source Data 4. Error bars on scatter plots show s.e.m.

**Figure S6-3.**
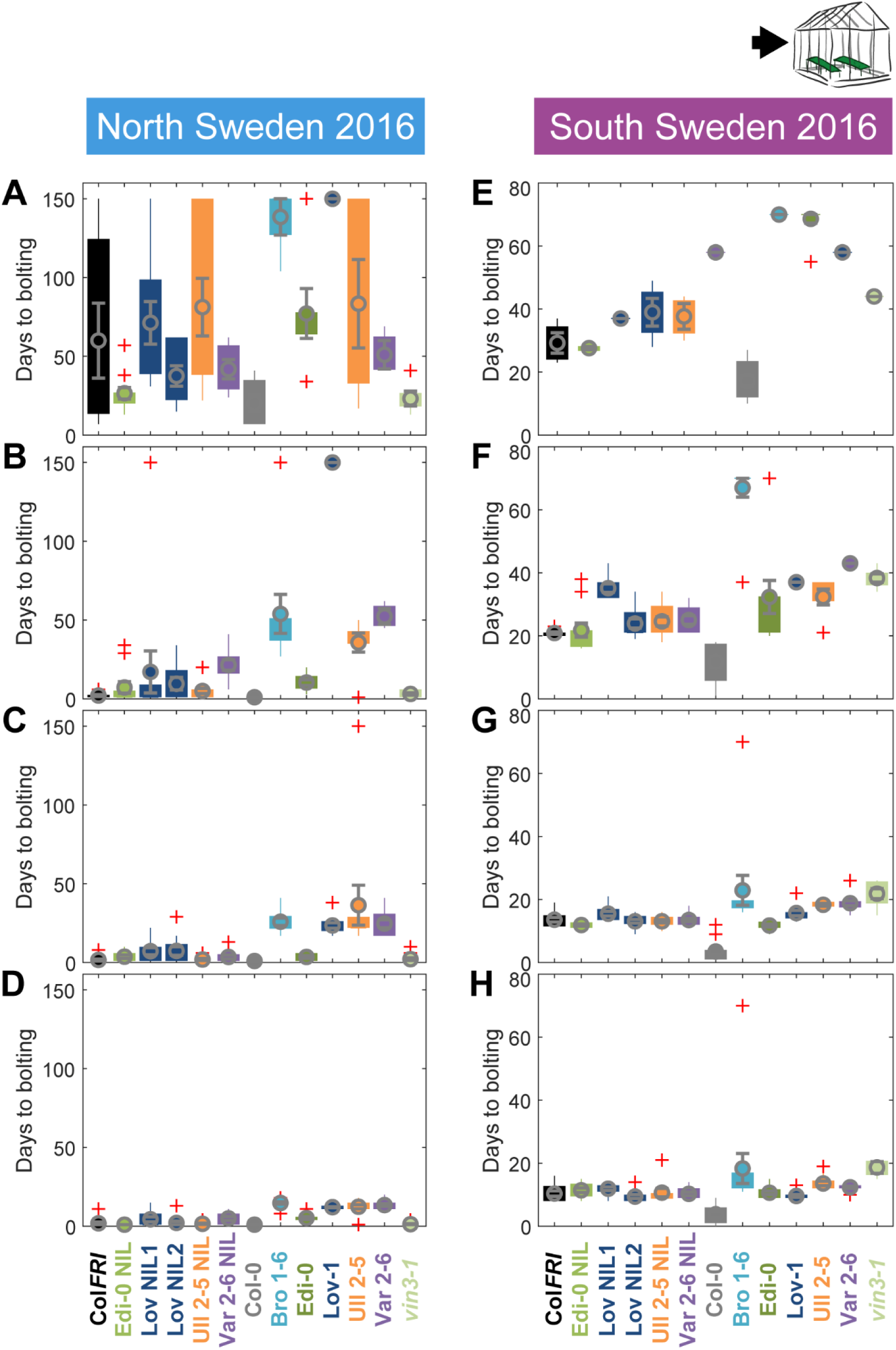
Bolting after transfer to warm, long-day conditions from winter in the field 2016-7 saturates at different rates in different genotypes in Sweden. Bolting time from sequential transfers to long day warm conditions from the field, for each genotype and transfer, n=12. **(A-D)** North Sweden, 06/09/2016, 04/10/2016, 01/11/2016,24/11/2016. **(E-H)** South Sweden, 01/10/2016, 22/10/2016, 19/11/2016, 17/12/2016. N=12, except where plants died, Source Data 5.

**Figure S6-4.**
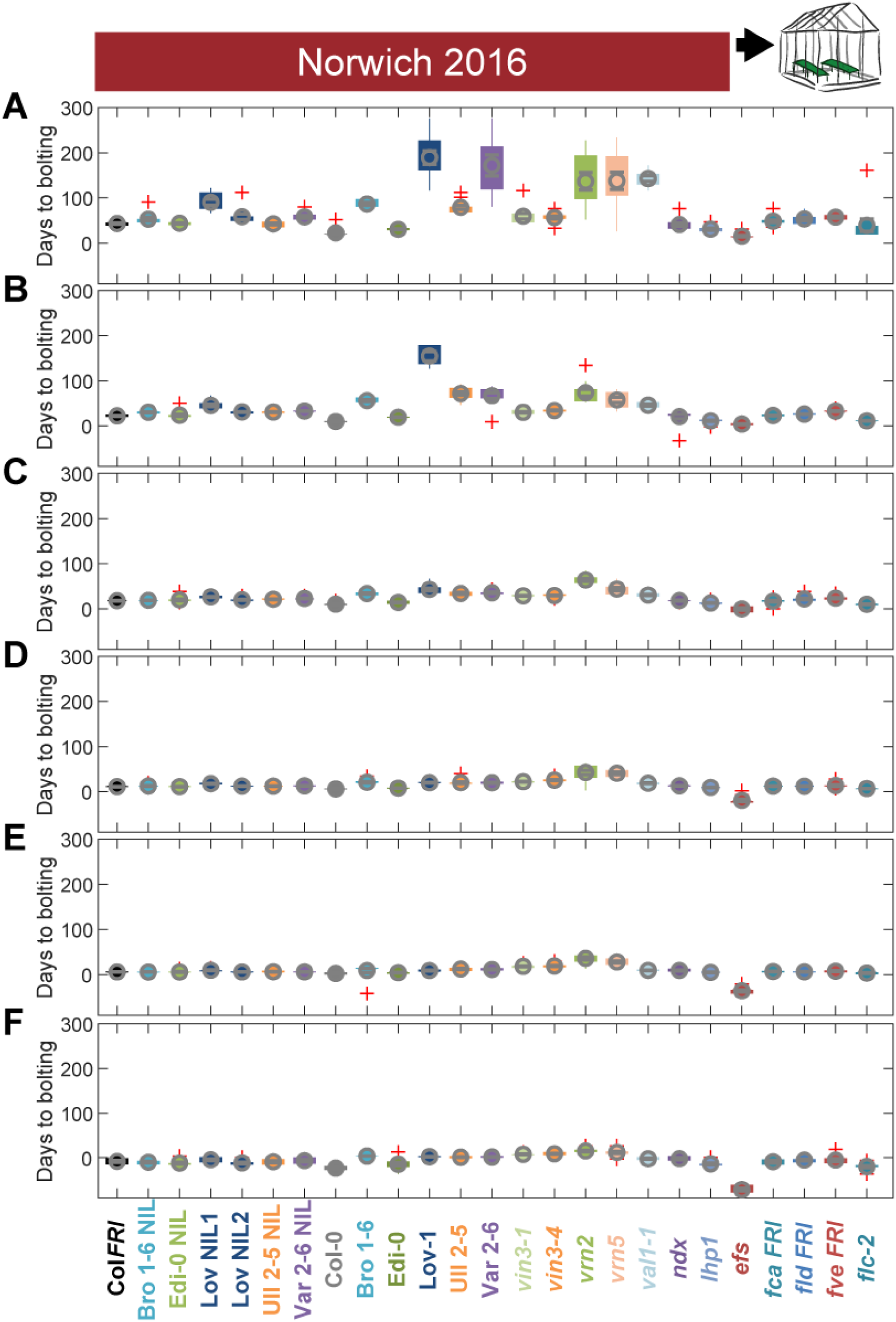
Bolting after transfer to warm, long-day conditions from winter in the field 2016-7 saturates at different rates in Norwich. Bolting from sequential transfers to long day warm conditions from the field, for each genotype and transfer, n=12. **(A)** 21/10/2016, **(B)** 03/11/2016, **(C)** 17/11/2016, **(D)** 30/11/2016, **(E)** 21/12/2016, **(F)** 26/01/2017. N=12, except where plants died, Source Data 5.

**Figure S6-5.**
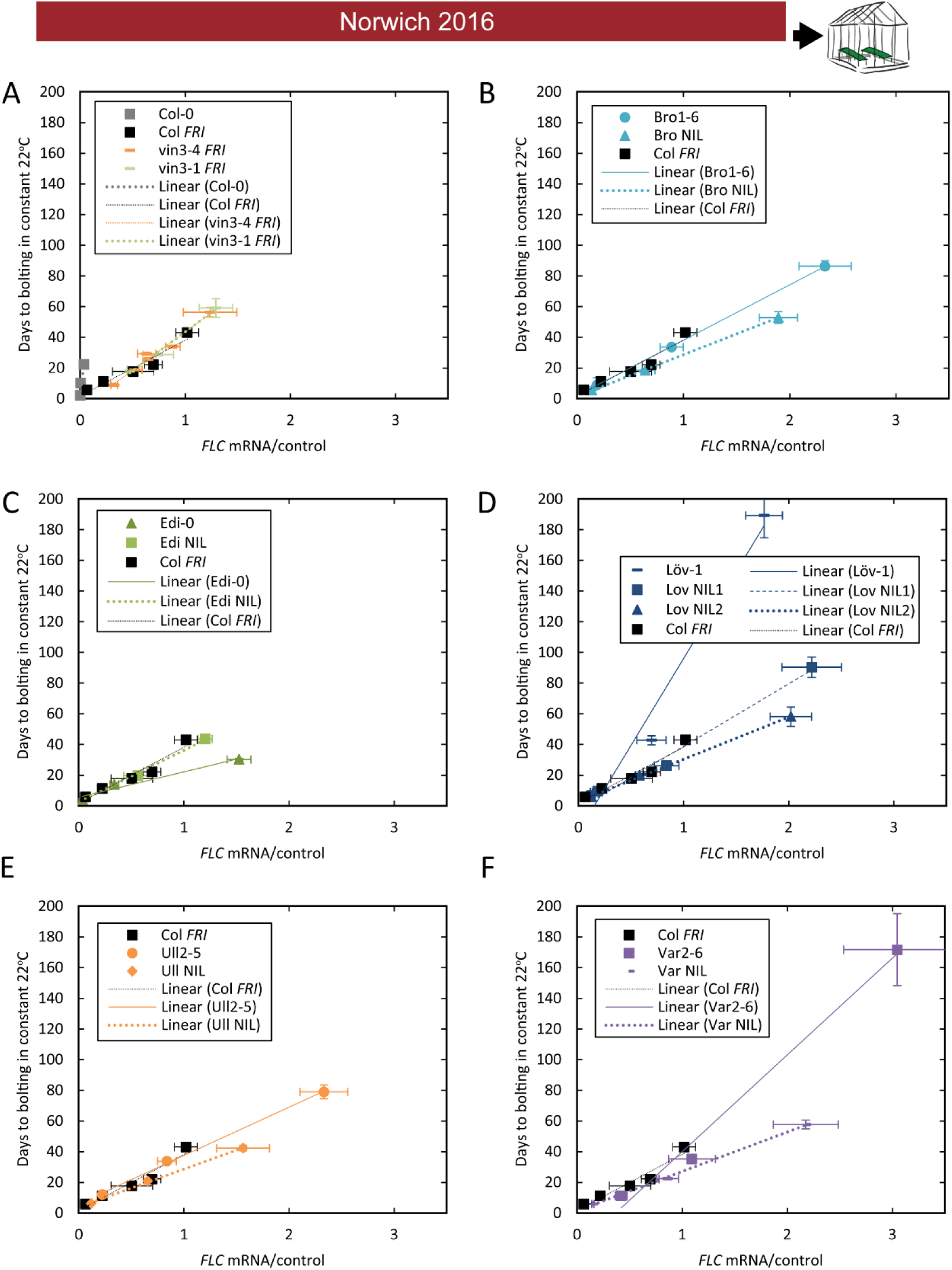
The relationship between time to floral transition and *FLC* expression at the end of cold (Norwich winter 2016-7) varies among accessions, both due to *trans* effects and due to the *FLC* alleles themselves. (**A-F**) Mean time to bolting of plants moved to a greenhouse lit for 16 hours, and maintained at 22°C/18°C light/dark, plotted against the mean *FLC* mRNA expression from plants sampled in the Norwich field condition greenhouse on the day of transfer, with linear regression lines plotted. For all accessions and NILs over 3-6 transfers at different times during the winter, R^2^= 0.68 for linear regression, p<0.001. N=6 for expression data, N=12 for bolting data, except where plants died, Source Data 3 and 5. Error bars show s.e.m.

**Figure S6-6.**
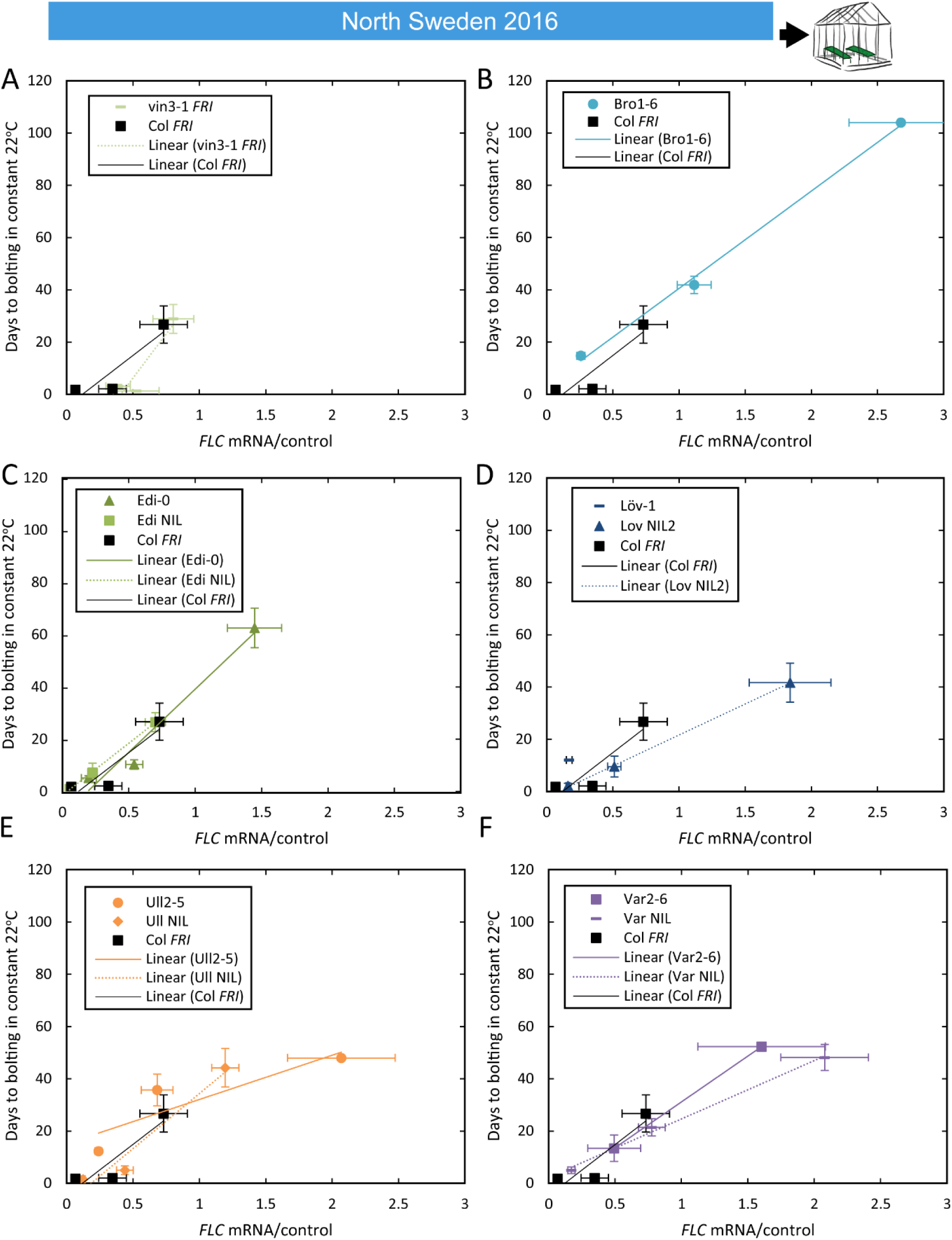
The relationship between time to floral transition and *FLC* expression at the end of cold in North Sweden winter 2016-7. Mean time to bolting of plants moved to a greenhouse lit for 16 hours, and maintained at 22°C, plotted against the mean *FLC* mRNA expression from plants sampled in the North Sweden field on or adjacent to the day of transfer, with linear regression lines plotted. (A-F) For all accessions and NILs over 3-6 transfers at different times during the winter, R^2^= 0.68 for linear regression, p<0.001. For D, there is no regression for Löv-1 as no Löv-1 plants from the first two transfers flowered within the 120 days of the experiment. N=12 for bolting data, except where plants died, Source Data 3 and 5. Error bars show s.e.m.

**Figure S7-1.**
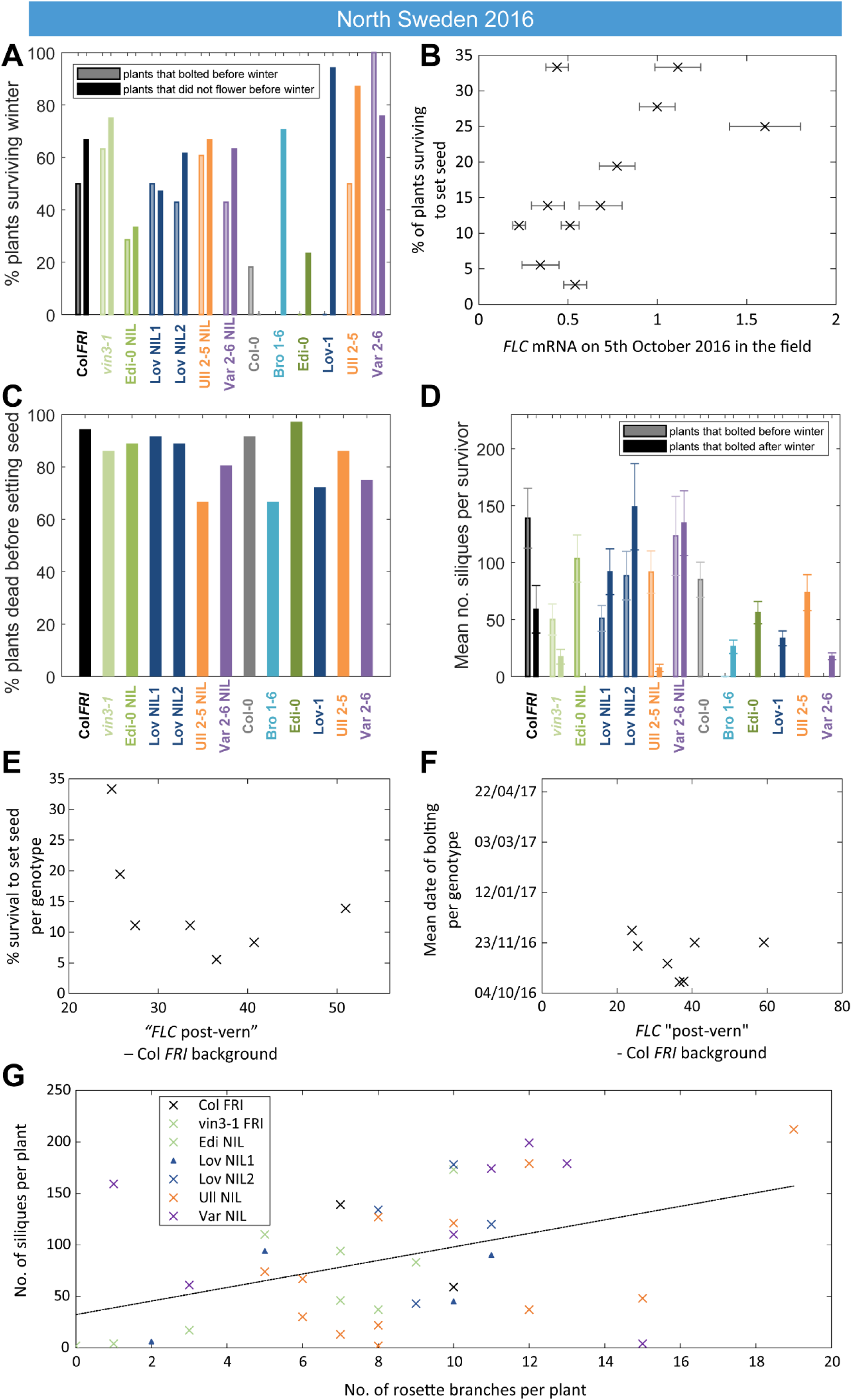
*FLC* controls fitness in North Sweden through bolting time and branching. Survival, branching and silique set in North Sweden are all correlated to aspects of *FLC* regulation. (**A**) Survival over winter of plants that bolted before winter in different genotypes versus survival of plants that did not bolt before winter. (**B**) Survival to seed set plotted against *FLC* levels (normalised to control sample for 2016-7) in the field in North Sweden 2016 (p<0.003, GLM for binomial data). (**C**) Percentage mortality before setting seed was high for all genotypes. (**D**) Mean number of siliques for plants surviving to set seed that bolted before or after winter. (**E**) Survival in the field does not correlate with *FLC* post-vern for the Col *FRI* background (GLM with binomial distribution, p-value > 0.1). (**F**) Date of bolting in the field does not correlate with *FLC* post-vern for the Col *FRI* background (linear regression, p-value > 0.1). (**G**) Silique production by surviving Col *FRI* background plants correlates with number of rosette branches, though more weakly at the individual level than at the genotype average level (linear regression, R^2^ = 0.23, p-value 0.004). N=36 plants sown (A, B, C, E) subsequent data based on survivors to seed set (D, G) and plants that survive to bolting (F), see Source Data 6.

**Figure S7-2.**
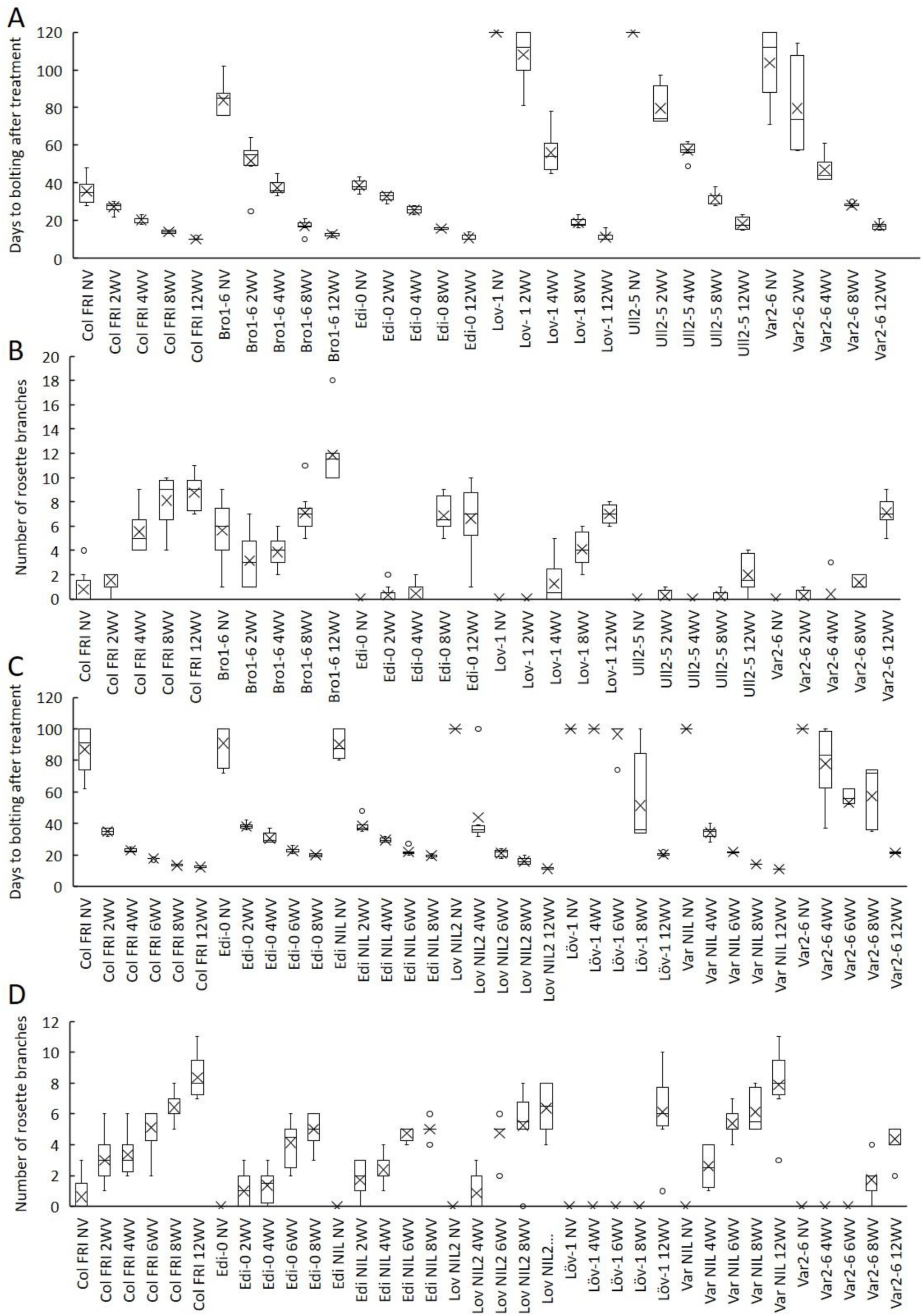
Increased vernalization increases the amount of rosette branch production and reduces the variability of bolting time. **(A)** Time to bolting for accessions in a heated, lit greenhouse without vernalization (NV) or after weeks of vernalization at constant 5°C (*n*WV), experiment 1. **(B)** Number of rosette branches for plants shown in A, experiment 1. **(C)** Time to bolting for selected accessions and NILs in a heated, lit greenhouse without vernalization (NV) or after weeks of vernalization at constant 5°C (*n*WV), experiment 2. **(D)** Number of rosette branches for plants shown in C, experiment 2. Median (central line), mean (cross), interquartile range (box), range (whiskers) and outliers (circles, values more than 1.5 times the interquartile range outside of the interquartile range). Plants that did not flower within 120 days of transfer not shown.

**Figure S7-3.**
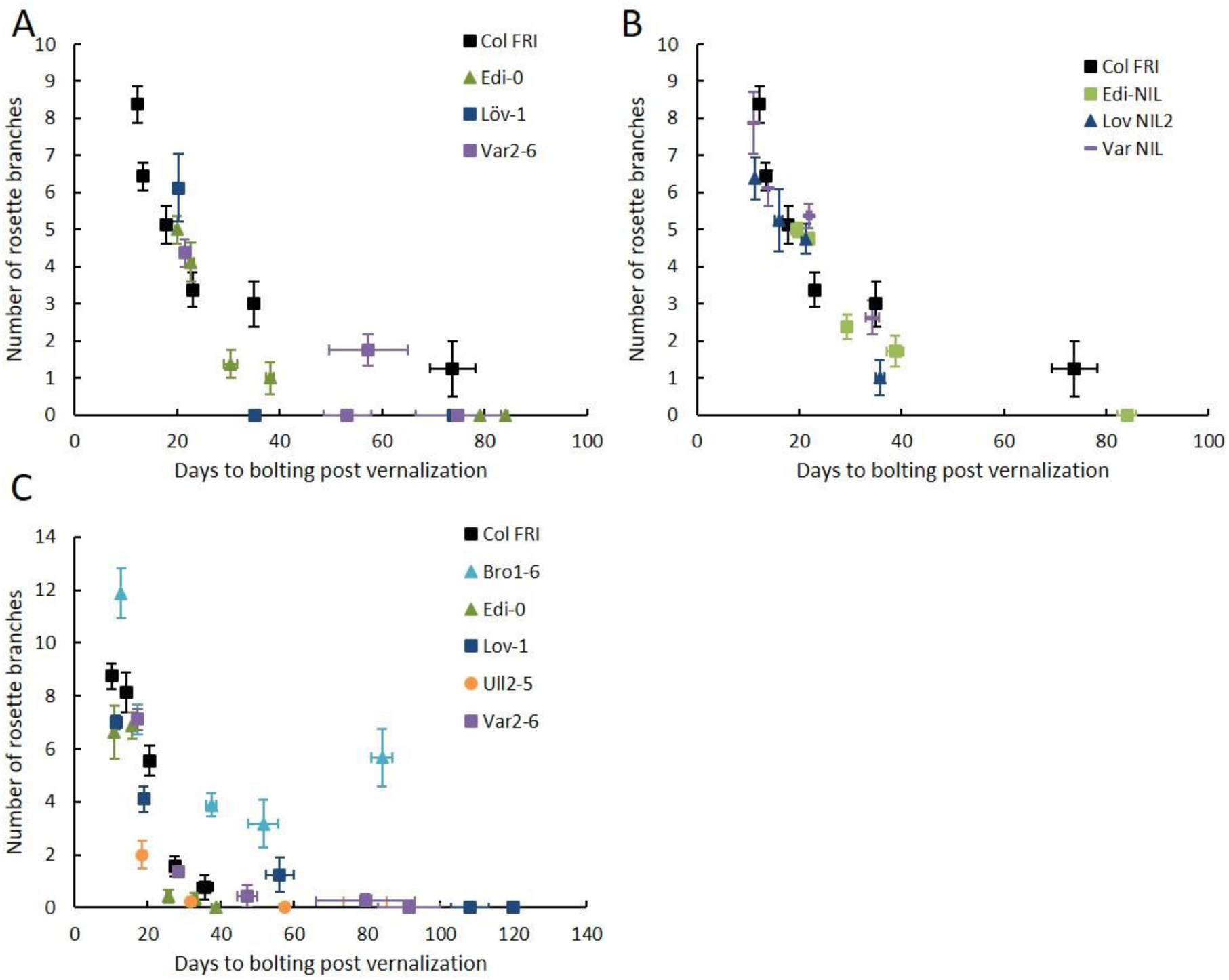
Increasing vernalization correlated with greater branch production with subtly different effects depending on *FLC* haplotype in the Col *FRI* background. Means per genotype and vernalization length treatment of rosette branch data presented in S7-2, plotted against days to bolting. (A) Data from S7-2A-B, experiment 1, accessions only. (B) Data from S7-2A-B, experiment 1, NILs only. (C) Data from S7-2C-D, experiment 2. Error bars show s.e.m.

**Figure S7-4.**
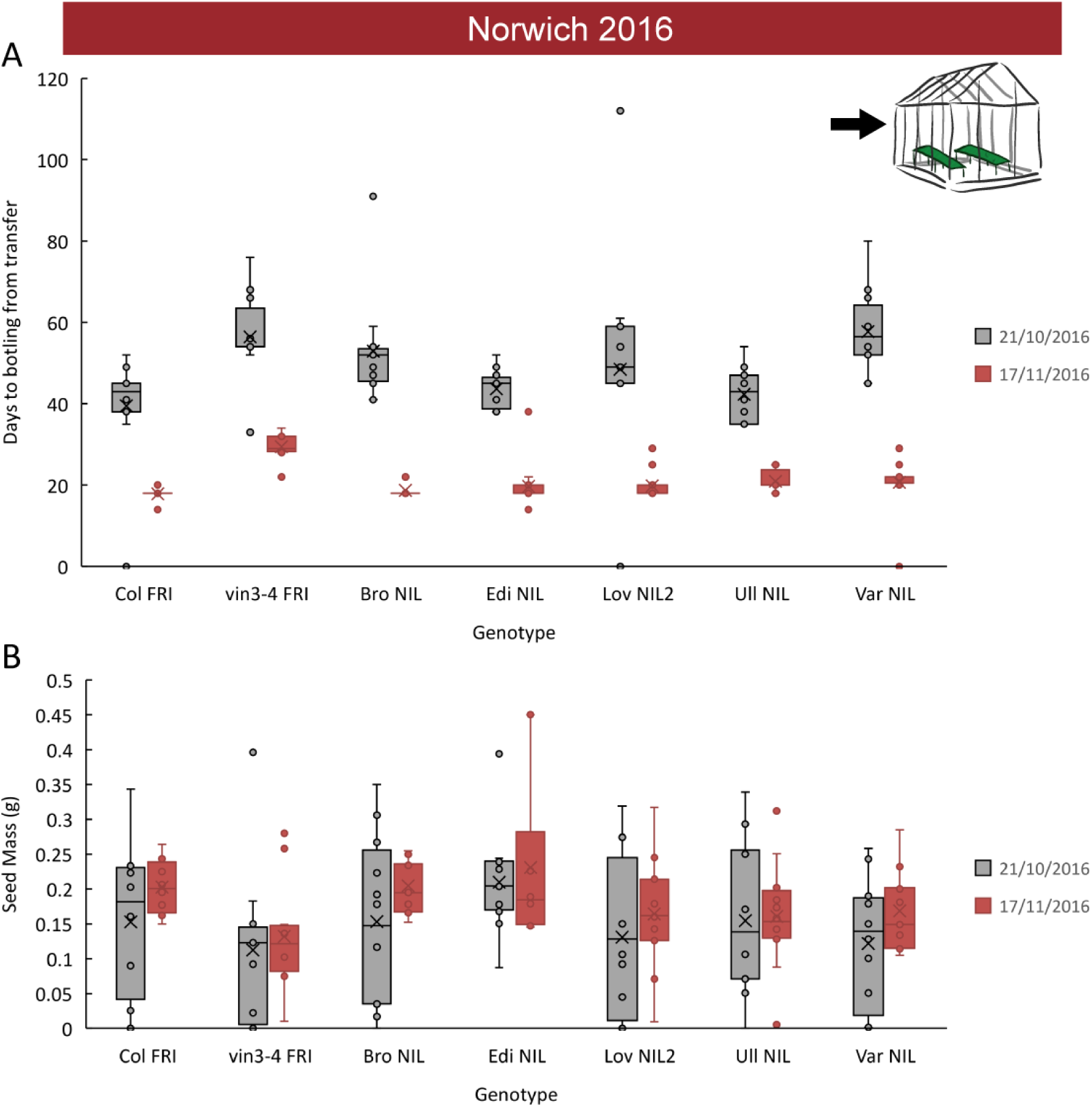
Increased vernalization increases the amount and reduces the variability of seed set. Flowering time for Col *FRI*, NILs and the *vin3-4 FRI* mutant after transfer to floral-induction conditions from ‘natural’ winter in Norwich 2016-7 on 21/10/16 and 17/11/16 (as in Fig. S6-4A, B). Total seed mass produced by plants in A. Box-and-whiskers plot for time to bolting for each genotype, showing median (central line), mean (cross), interquartile range (box), range (whiskers) and outliers (circles, values more than 1.5 times the interquartile range outside of the interquartile range). The time to bolt per plant negatively correlated with seed mass produced, p<0.001, Kenward-Roger’s t-test on REML Linear mixed model with date of transfer as a random factor.

## References

Ågren J, Oakley CG, Lundemo S, Schemske DW (2016) Adaptive divergence in flowering time among natural populations of Arabidopsis thaliana: Estimates of selection and QTL mapping. Evolution n/a-n/a

Ågren J, Oakley CG, McKay JK, Lovell JT, Schemske DW (2013) Genetic mapping of adaptation reveals fitness tradeoffs in Arabidopsis thaliana. Proc Natl Acad Sci 110: 21077–21082

Andrés F, Coupland G (2012) The genetic basis of flowering responses to seasonal cues. Nat Rev Genet 13: 627–639

Angel A, Song J, Dean C, Howard M (2011) A Polycomb-based switch underlying quantitative epigenetic memory. Nature 476: 105–108

Antoniou-Kourounioti RL, Hepworth J, Heckmann A, Duncan S, Qüesta J, Rosa S, Säll T, Holm S, Dean C, Howard M (2018) Temperature Sensing Is Distributed throughout the Regulatory Network that Controls FLC Epigenetic Silencing in Vernalization. Cell Syst 7: 643-655.e9

Auge GA, Penfield S, Donohue K (2019) Pleiotropy in developmental regulation by flowering-pathway genes: is it an evolutionary constraint? New Phytol 224: 55–70

Ausín I, Alonso-Blanco C, Jarillo JA, Ruiz-García L, Martínez-Zapater JM (2004) Regulation of flowering time by FVE, a retinoblastoma-associated protein. Nat Genet 36: 162–166

Berry S, Hartley M, Olsson TSG, Dean C, Howard M (2015) Local chromatin environment of a Polycomb target gene instructs its own epigenetic inheritance. eLife e07205

Bloomer RH, Dean C (2017) Fine-tuning timing: natural variation informs the mechanistic basis of the switch to flowering in Arabidopsis thaliana. J Exp Bot 68: 5439–5452

Bond DM, Wilson IW, Dennis ES, Pogson BJ, Jean Finnegan E (2009) VERNALIZATION INSENSITIVE 3 (VIN3) is required for the response of Arabidopsis thaliana seedlings exposed to low oxygen conditions. Plant J 59: 576–587

Burghardt LT, Runcie DE, Wilczek AM, Cooper MD, Roe JL, Welch SM, Schmitt J (2016) Fluctuating, warm temperatures decrease the effect of a key floral repressor on flowering time in Arabidopsis thaliana. New Phytol 210: 564–576

Coustham V, Li P, Strange A, Lister C, Song J, Dean C (2012) Quantitative modulation of polycomb silencing underlies natural variation in vernalization. Science 337: 584–587

Czechowski T, Stitt M, Altmann T, Udvardi MK, Scheible W-R (2005) Genome-Wide Identification and Testing of Superior Reference Genes for Transcript Normalization in Arabidopsis. Plant Physiol 139: 5–17

De Lucia F, Crevillen P, Jones AME, Greb T, Dean C (2008) A PHD-polycomb repressive complex 2 triggers the epigenetic silencing of FLC during vernalization. Proc Natl Acad Sci U S A 105: 16831–16836

Dittmar EL, Oakley CG, Ågren J, Schemske DW (2014) Flowering time QTL in natural populations of Arabidopsis thaliana and implications for their adaptive value. Mol Ecol 23: 4291–4303

Duncan S, Holm S, Questa J, Irwin J, Grant A, Dean C (2015) Seasonal shift in timing of vernalization as an adaptation to extreme winter. eLife 4: e06620

Feltz CJ, Miller GE (1996) An asymptotic test for the equality of coefficients of variation from k populations. Stat Med 15: 646–658

Fournier-Level A, Korte A, Cooper MD, Nordborg M, Schmitt J, Wilczek AM (2011) A map of local adaptation in Arabidopsis thaliana. Science 334: 86–89

Fournier-Level A, Perry EO, Wang JA, Braun PT, Migneault A, Cooper MD, Metcalf CJE, Schmitt J (2016) Predicting the evolutionary dynamics of seasonal adaptation to novel climates in Arabidopsis thaliana. Proc Natl Acad Sci U S A 113: E2812–2821

Greb T, Mylne JS, Crevillen P, Geraldo N, An H, Gendall AR, Dean C (2007) The PHD finger protein VRN5 functions in the epigenetic silencing of Arabidopsis FLC. Curr Biol CB 17: 73–78

Grillo MA, Li C, Hammond M, Wang L, Schemske DW (2013) Genetic architecture of flowering time differentiation between locally adapted populations of Arabidopsis thaliana. New Phytol 197: 1321–1331

Hepworth J, Antoniou-Kourounioti RL, Bloomer RH, Selga C, Berggren K, Cox D, Collier Harris BR, Irwin JA, Holm S, Säll T, et al (2018) Absence of warmth permits epigenetic memory of winter in Arabidopsis. Nat Commun 9: 639

Huang X, Ding J, Effgen S, Turck F, Koornneef M (2013) Multiple loci and genetic interactions involving flowering time genes regulate stem branching among natural variants of Arabidopsis. New Phytol 199: 843–857

Jong M de, Tavares H, Pasam RK, Butler R, Ward S, George G, Melnyk CW, Challis R, Kover PX, Leyser O (2019) Natural variation in Arabidopsis shoot branching plasticity in response to nitrate supply affects fitness. PLOS Genet 15: e1008366

Jung J-H, Park J-H, Lee S, To TK, Kim J-M, Seki M, Park C-M (2013) The cold signaling attenuator HIGH EXPRESSION OF OSMOTICALLY RESPONSIVE GENE1 activates FLOWERING LOCUS C transcription via chromatin remodeling under short-term cold stress in Arabidopsis. Plant Cell 25: 4378–4390

Kim H-J, Hyun Y, Park J-Y, Park M-J, Park M-K, Kim MD, Kim H-J, Lee MH, Moon J, Lee I, et al (2004) A genetic link between cold responses and flowering time through FVE in Arabidopsis thaliana. Nat Genet 36: 167–171

Kudoh H (2016) Molecular phenology in plants: in natura systems biology for the comprehensive understanding of seasonal responses under natural environments. New Phytol 210: 399–412

Lazaro A, Obeng-Hinneh E, Albani MC (2018) Extended Vernalization Regulates Inflorescence Fate in Arabis alpina by Stably Silencing PERPETUAL FLOWERING1. Plant Physiol 176: 2819–2833

Lee I, Amasino RM (1995) Effect of Vernalization, Photoperiod, and Light Quality on the Flowering Phenotype of Arabidopsis Plants Containing the FRIGIDA Gene. Plant Physiol 108: 157–162

Lempe J, Balasubramanian S, Sureshkumar S, Singh A, Schmid M, Weigel D (2005) Diversity of Flowering Responses in Wild Arabidopsis thaliana Strains. PLoS Genet 1: e6

Li P, Filiault D, Box MS, Kerdaffrec E, van Oosterhout C, Wilczek AM, Schmitt J, McMullan M, Bergelson J, Nordborg M, et al (2014) Multiple FLC haplotypes defined by independent cis-regulatory variation underpin life history diversity in Arabidopsis thaliana. Genes Dev 28: 1635–1640

Li P, Tao Z, Dean C (2015) Phenotypic evolution through variation in splicing of the noncoding RNA COOLAIR. Genes Dev. doi: 10.1101/gad.258814.115

Liu F, Marquardt S, Lister C, Swiezewski S, Dean C (2010) Targeted 3′ Processing of Antisense Transcripts Triggers Arabidopsis FLC Chromatin Silencing. Science 327: 94–97

Liu F, Quesada V, Crevillén P, Bäurle I, Swiezewski S, Dean C (2007) The Arabidopsis RNA-binding protein FCA requires a lysine-specific demethylase 1 homolog to downregulate FLC. Mol Cell 28: 398–407

Marwick B, Krishnamoorthy K (2019) cvequality: Tests for the equality of coefficients of variation from multiple groups. R software package version 0.1.3. Retrieved from github.com/benmarwick/cvequality on 07/01/2019.

Méndez-Vigo B, Picó FX, Ramiro M, Martínez-Zapater JM, Alonso-Blanco C (2011) Altitudinal and climatic adaptation is mediated by flowering traits and FRI, FLC, and PHYC genes in Arabidopsis. Plant Physiol 157: 1942–1955

Michaels SD, Amasino RM (1999) FLOWERING LOCUS C Encodes a Novel MADS Domain Protein That Acts as a Repressor of Flowering. Plant Cell Online 11: 949–956

Mylne JS, Barrett L, Tessadori F, Mesnage S, Johnson L, Bernatavichute YV, Jacobsen SE, Fransz P, Dean C (2006) LHP1, the Arabidopsis homologue of HETEROCHROMATIN PROTEIN1, is required for epigenetic silencing of FLC. Proc Natl Acad Sci U S A 103: 5012–5017

Nagano AJ, Kawagoe T, Sugisaka J, Honjo MN, Iwayama K, Kudoh H (2019) Annual transcriptome dynamics in natural environments reveals plant seasonal adaptation. Nat Plants 5: 74–83

Nagel DH, Doherty CJ, Pruneda-Paz JL, Schmitz RJ, Ecker JR, Kay SA (2015) Genome-wide identification of CCA1 targets uncovers an expanded clock network in Arabidopsis. Proc Natl Acad Sci 112: E4802–E4810

Qüesta JI, Antoniou-Kourounioti RL, Rosa S, Li P, Duncan S, Whittaker C, Howard M, Dean C (2020) Noncoding SNPs influence a distinct phase of Polycomb silencing to destabilize long-term epigenetic memory at Arabidopsis FLC. Genes Dev 34: 446–461

Qüesta JI, Song J, Geraldo N, An H, Dean C (2016) Arabidopsis transcriptional repressor VAL1 triggers Polycomb silencing at FLC during vernalization. Science 353: 485–488

R Core Team (2018) R: A Language and Environment for Statistical Computing. R Foundation for Statistical Computing, Vienna, Austria

Rubin MJ, Brock MT, Baker RL, Wilcox S, Anderson K, Davis SJ, Weinig C (2018) Circadian rhythms are associated with shoot architecture in natural settings. New Phytol 219: 246–258

Ruijter JM, Ramakers C, Hoogaars WMH, Karlen Y, Bakker O, Hoff VD, B MJ, Moorman AFM (2009) Amplification efficiency: linking baseline and bias in the analysis of quantitative PCR data. Nucleic Acids Res 37: e45–e45

Sánchez-Bermejo E, Méndez-Vigo B, Picó FX, Martínez-Zapater JM, Alonso-Blanco C (2012) Novel natural alleles at FLC and LVR loci account for enhanced vernalization responses in Arabidopsis thaliana: FLC and LVR increase vernalization responses. Plant Cell Environ 35: 1672–1684

Sasaki E, Frommlet F, Nordborg M (2018) GWAS with Heterogeneous Data: Estimating the Fraction of Phenotypic Variation Mediated by Gene Expression Data. G3 Genes Genomes Genet 8: 3059–3068

Sasaki E, Zhang P, Atwell S, Meng D, Nordborg M (2015) “Missing” G x E Variation Controls Flowering Time in Arabidopsis thaliana. PLoS Genet 11: e1005597

Sheldon CC, Burn JE, Perez PP, Metzger J, Edwards JA, Peacock WJ, Dennis ES (1999) The FLF MADS box gene: a repressor of flowering in Arabidopsis regulated by vernalization and methylation. Plant Cell 11: 445–458

Shindo C, Lister C, Crevillen P, Nordborg M, Dean C (2006) Variation in the epigenetic silencing of FLC contributes to natural variation in Arabidopsis vernalization response. Genes Dev 20: 3079–3083

Song YH, Kubota A, Kwon MS, Covington MF, Lee N, Taagen ER, Cintrón DL, Hwang DY, Akiyama R, Hodge SK, et al (2018) Molecular basis of flowering under natural long-day conditions in Arabidopsis. Nat Plants 4: 824–835

Strange A, Li P, Lister C, Anderson J, Warthmann N, Shindo C, Irwin J, Nordborg M, Dean C (2011) Major-effect alleles at relatively few loci underlie distinct vernalization and flowering variation in Arabidopsis accessions. PloS One 6: e19949

Sun Q, Csorba T, Skourti-Stathaki K, Proudfoot NJ, Dean C (2013) R-Loop Stabilization Represses Antisense Transcription at the Arabidopsis FLC Locus. Science 340: 619–621

Sung S, Amasino RM (2004) Vernalization in Arabidopsis thaliana is mediated by the PHD finger protein VIN3. Nature 427: 159–164

Taylor MA, Wilczek AM, Roe JL, Welch SM, Runcie DE, Cooper MD, Schmitt J (2019) Large-effect flowering time mutations reveal conditionally adaptive paths through fitness landscapes in Arabidopsis thaliana. Proc Natl Acad Sci 116: 17890–17899

Urbaniak GC, Plous S (2015) Research Randomizer (Version 4.0).

Wang R, Farrona S, Vincent C, Joecker A, Schoof H, Turck F, Alonso-Blanco C, Coupland G, Albani MC (2009) PEP1 regulates perennial flowering in Arabis alpina. Nature 459: 423–427

Wilczek AM, Roe JL, Knapp MC, Cooper MD, Lopez-Gallego C, Martin LJ, Muir CD, Sim S, Walker A, Anderson J, et al (2009) Effects of Genetic Perturbation on Seasonal Life History Plasticity. Science 323: 930–934

Wu Z, Ietswaart R, Liu F, Yang H, Howard M, Dean C (2016) Quantitative regulation of FLC via coordinated transcriptional initiation and elongation. Proc Natl Acad Sci 113: 218–223

Yang H, Berry S, Olsson TSG, Hartley M, Howard M, Dean C (2017) Distinct phases of Polycomb silencing to hold epigenetic memory of cold in Arabidopsis. Science 357: 1142–1145

Yang H, Howard M, Dean C (2014) Antagonistic Roles for H3K36me3 and H3K27me3 in the Cold-Induced Epigenetic Switch at Arabidopsis FLC. Curr Biol 24: 1793–1797

Yuan W, Luo X, Li Z, Yang W, Wang Y, Liu R, Du J, He Y (2016) A *cis* cold memory element and a *trans* epigenome reader mediate Polycomb silencing of *FLC* by vernalization in *Arabidopsis*. Nat Genet 48: 1527–1534

